# Nickel-NTA lipid-monolayer affinity grids allow for high-resolution structure determination by cryo-EM

**DOI:** 10.1101/2025.04.24.650495

**Authors:** Aleksandra Skrajna, Emily Robinson, Kevin Cannon, Reta Sarsam, Richard G. Ouellette, Patrick Brennwald, Robert K. McGinty, Joshua D. Strauss, Richard W. Baker

**Affiliations:** Center for Integrative Chemical Biology and Drug Discovery, Division of Chemical Biology and Medicinal Chemistry, UNC Eshelman School of Pharmacy; Chapel Hill, NC, USA; Department of Biochemistry and Biophysics, UNC-Chapel Hill School of Medicine; Chapel Hill, NC, USA; Department of Cell Biology and Physiology, UNC Chapel Hill School of Medicine; Chapel Hill, NC, USA

## Abstract

Grid preparation is a rate-limiting step in determining high-resolution structures by single particle cryogenic electron microscopy (cryo-EM). Particle interaction with the air-water interface often leads to denaturation, aggregation, or a preferred orientation within the ice. Some samples yield insufficient quantities of particles when using traditional grid making techniques and require the use of solid supports that concentrate samples onto the grid. Recent advances in grid-preparation show that affinity grids are promising tools to selectively concentrate proteins while simultaneously protecting samples from the air-water interface. One such technique utilizes lipid monolayers containing a lipid species with an affinity handle. Some of the first affinity grids used a holey carbon layer coated with nickel nitrilotriacetic acid (Ni-NTA) lipid, which allowed for the binding of proteins bearing the commonly used poly-histidine affinity tag. These studies however used complicated protocols and were conducted before the “resolution revolution” of cryo-EM. Here, we provide a straight-forward preparation method and systematic analysis of Ni-NTA lipid monolayers as a tool for high-resolution single particle cryo-EM. We found that lipid affinity grids concentrate particles away from the air-water interface in thin ice (∼30 nm). We determined a 2.6 Å structure of the human nucleosome, showing this method is amenable to high-resolution structure determination. Furthermore, we determined a 3.1 Å structure of a sub-100 kDa protein demonstrating that this technique is amenable to proteins across biological size ranges. Lipid monolayers are therefore an easily extendable tool for most systems and help alleviate common problems such as low yield, disruption by the air-water interface, and thicker ice.

## Introduction

Single-particle electron cryo-microscopy (cryo-EM) is a dominant technique in the field of structural biology, with the so-called “resolution revolution” of the past decade cementing its place as a go-to method for high-resolution structure determination of biological macromolecules [1]. Comparatively speaking, sample preparation methods have remained relatively unchanged, with the vast majority of deposited structures prepared using traditional blotting methods that suspend a thin layer of aqueous sample within the open holes of a cryo-EM grid. While many proteins are amenable to traditional and straight-forward methods, sample preparation and grid making are still rate-limiting steps in the structure determination pipeline, and many samples ultimately fail to produce structures or require an immense amount of optimization. The “ideal” cryo-grid contains areas of thin vitreous ice, densely populated with a single layer of randomly oriented particles. The most common issue at the specimen level comes as a result of the pernicious “air-water interface” (AWI), which often results in protein denaturation, preferred orientation, low particle number, or too high particle number with overlapping layers of particles at both AWIs [2, 3]. Non-traditional grid preparation methods largely revolve around solving issues introduced by the AWI [4-6].

A variety of solid supports have been developed to coat cryo-EM grids, used as non-selective surfaces that concentrate samples on the grid and partially protect from the AWI. Perhaps the earliest and most commonly used grid support is thin amorphous carbon, which has led to a variety of highresolution reconstructions [7-10]. As such, carbon evaporators are a common piece of equipment in most cryo-EM facilities. More recently, many labs have developed the use of graphene or graphene oxide, which theoretically provides many of the benefits of amorphous carbon while contributing far less background signal due to the 1-atom thickness of graphene [11]. However, graphene itself is highly hydrophobic and must be functionalized before use, either through chemical modification or the use of a UV ozone cleaner [12]. While graphene requires a multi-step process to coat the grid with a single 2D crystal, graphene oxide is conveniently available as “flakes” that can be applied to grids, although this process is prone to poor coverage, multiple overlapping layers, and a lack of reproducibility [13]. Despite these challenges, several groups have reported highresolution structures in which graphene supports were used to troubleshoot a difficult sample [9, 14-16].

Similar to carbon and graphene, the use of affinity grids provides many of the benefits of solid supports, while also providing a selectivity filter to deposit samples from a complicated milieu (e.g. cell extracts) or to concentrate low abundance samples. Consisting of a chemically modified TEM grid and designed to bind selectively to a target protein complex, these grids have proven useful in determining the structure of samples not amenable to traditional TEM grid design or expanding the cryo-EM toolkit to novel methods like on-grid purification. Among these affinity grids include chemically-modified carbon films [17, 18], chemically-modified graphene oxide [15, 19], 2D crystals of streptavidin designed to bind biotin-conjugated samples [20, 21], and lipid monolayers with functionalized polar head groups [22, 23].

One promising and potentially broad-reaching grid support is the use of nickel nitrilotriacetic acid (Ni-NTA) lipid monolayers to bind proteins bearing a poly-histidine purification tag, the most common tag for recombinant purification of proteins [24]. Ni-NTA monolayers have been used to concentrate large protein complexes and viruses onto the TEM grid for both cryo-EM and negative stain applications [25-29], but these studies were conducted before the advent of direct-electron detectors and the “resolution revolution” of the past decade. It is therefore unclear whether grids with a Ni-NTA monolayer are amenable to high-resolution structure determination and what are the limitations and general guidelines for their use in the modern cryo-EM landscape. In this study we systematically explore the use of Ni-NTA affinity grids at our Multiuser Core Facility and provide general guidelines for extending this technology to other projects. We also provide a robust and easy-to-use protocol for preparing lipid monolayer grids and demonstrate its ease of use with novice users.

## Results

### Overview of Ni-NTA lipid monolayers as a grid support

Many projects ultimately fail to produce high-resolution structures at our multi-user core facility. The most common problems we observe are preferred orientation, aggregation or complex disassembly at the AWI, and proteins that do not enter the grid hole. We therefore set out to explore the utility and reproducibility of lipid monolayers as a grid support available to our users. We provided onsite training to 5 core-users (3 postdoctoral fellows, and two facility staff), all of which had previous experience operating a Vitrobot plunge-freezing device but did not have any prior experience preparing lipid monolayers. Our protocol for coating grids with lipid monolayers is based on previous methods for producing 2D-layers of streptavidin on a monolayer of biotin-conjugated lipid [4]. However, we have streamlined the process for use of lipid monolayers as a generalizable affinity support. The lipid monolayers are formed on a droplet of castor oil and deposited on a grid by gently touching the grid to the surface of the droplet (Figure 1A,B). Grids are then washed in a background buffer equivalent to the protein buffer, Histagged protein is incubated for a variable time, followed by additional buffer washes, and plunge-freezing on a Vitrobot Mark IV plunge freezing device. We designed a 3D-printed manifold that holds up to 8 pairs of forceps in a home-made humidity chamber or a Vitrobot Mark IV sample chamber. This maintains a high humidity (∼95%) during sample deposition, allowing us to prepare multiple grids in a single ∼2 hour freezing session.

**Figure 1.**
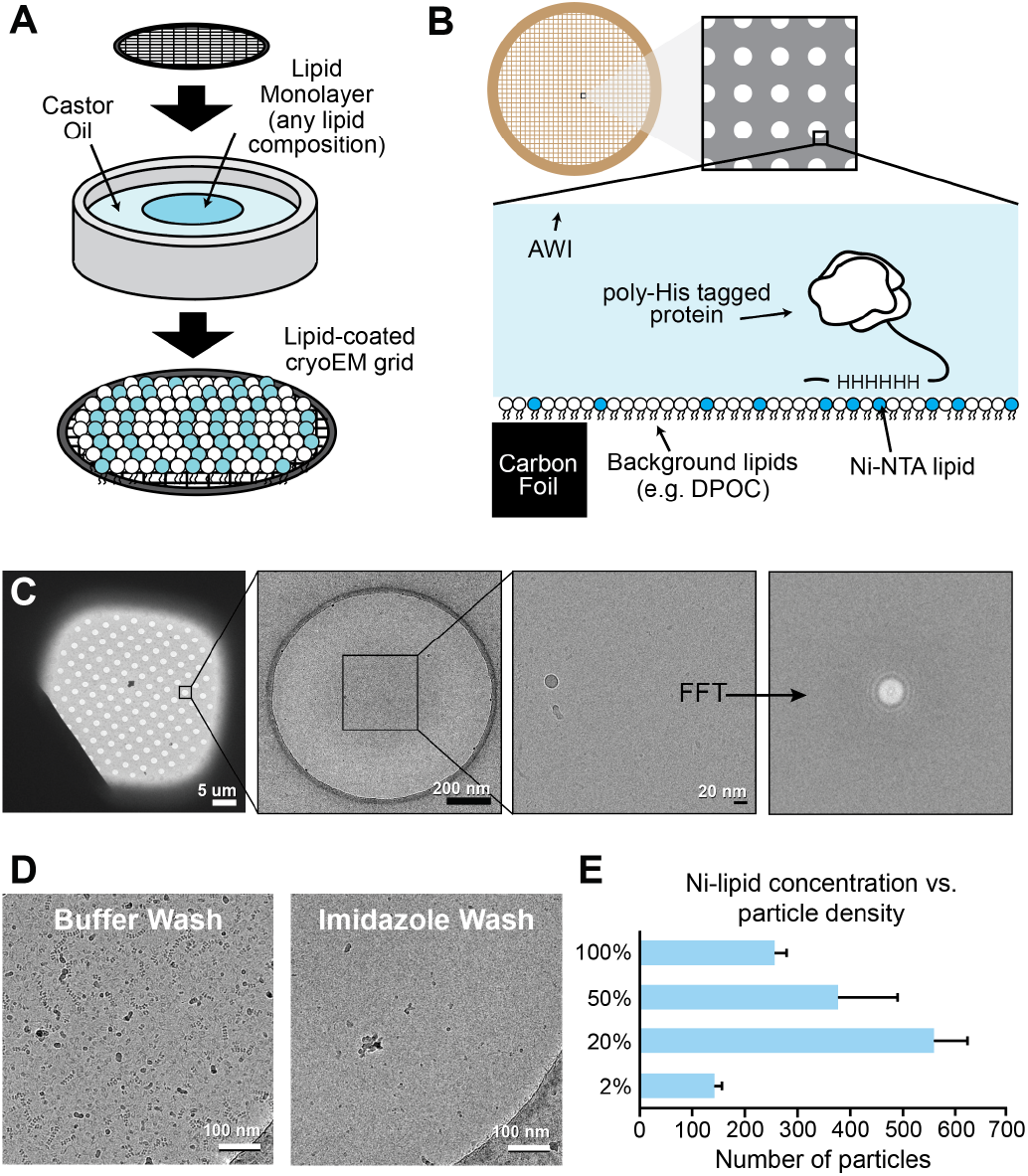
Design and implementation of lipid monolayer affinity grids. **(A)** Schematic for deposition of lipid monolayers on a cryo-EM grid. **(B)** Diagram of lipid monolayer on a cryo-EM grid, with position of carbon foil, lipid, protein, and air-water interface (AWI) noted. **(C)** Micrographs taken at different magnifications showing a lipid-coated grid at the square, hole, and record level. The FFT of a high magnification image is shown (far right). **(D)** Comparison of hNuc sample deposited on a Ni-NTA grid with a buffer wash (left) and an imidazole wash (right), which disrupts the interaction of the His-tag with the Ni-NTA lipid. **(E)** Quantification of particle number on Ni-NTA lipid grids containing a variable amount of Ni-NTA lipid.

### Characterization of Ni-NTA lipid monolayer grids

To begin our analysis, we made lipid monolayer grids with our standard protocol and screened grids prepared without protein. We observed robust coverage of our grids, denoted by minimal broken squares (Supp. Fig. 1A,B) and observable thon rings in the FFT of high-magnification images taken within ice-containing grid holes (Figure 1C). To demonstrate that the Ni-NTA lipid monolayer could recruit proteins in a His-tag-dependent manner, we made grids and incubated them with a nucleosome bearing a His-tag on the N-terminus of histone H2A (hereon referred to as hNuc). Ni-NTA grids were incubated with hNuc and then washed with a base buffer or a buffer containing 200 mM Imidazole. As shown in Figure 1D, the high imidazole wash results in no observable hNuc particles, indicating that binding is a direct result of chelation between the Ni-NTA moiety and the imidazole sidechains of the histidine residues.

We next sought to test the best lipid composition for our monolayer grids. While some other protocols for growing 2D crystals use 100% of a single lipid species to make monolayers [30-32], we supposed that the concentrated charge of the Ni-NTA moiety would make a lipid monolayer of 100% Ni-NTA lipid unstable. Furthermore, previous use of Ni-NTA lipids on carbon grids showed that concentrations as low as 2% are sufficient to recruit proteins [25]. We therefore tested a variety of Ni-NTA lipid concentrations (2, 20, 50, 100 %) in a background of DOPC, a lipid with a neutral head group. Each grid was made otherwise identically and incubated with hNuc at a concentration of 0.05 mg/mL. For each sample, we collected multiple images in grid squares containing roughly equivalent ice thickness, as judged by the number of visible holes in the SerialEM montage. Particles were manually picked in IMOD [33] and a per-micrograph average particle number was calculated. While all lipid concentrations contained visible particles (Supp. Fig. 1D), the 20% and 50% samples both contained the most particles (Figure 1E). Interestingly, the 100% Ni-NTA condition contained only 1/3 the number of particles of the lower concentrations suggesting issues with the formation of uniform lipid monolayer. Moving forward, we decided to use 20% Ni-NTA for fabrication of affinity grids

### Ni-NTA lipid monolayers concentrate particles away from the AWI onto the surface of the TEM grid

One benefit of affinity grids is the potential to concentrate low-abundance samples. Whereas traditional grid preparation methods typically require proteins at concentrations of at least 1 mg/mL, grids with less restrictive sample requirements might allow for imaging of “precious” samples that can only be isolated in very limited quantities, or allow for on-grid purification from just a few microliters of cell extract. To test the ability of Ni-NTA grids to concentrate His-tagged proteins compared to holey carbon grids, we prepared holey carbon and lipid monolayer grids for two samples: His-tagged glutamine synthase (GS) and hNuc. For each sample, we used a high-concentration for the standard grid prep, and a 20- or 40-fold lower protein concentration for the Ni-NTA affinity grid.

To determine orientation of particles in ice and measure the concentrating effect on the lipid film, we imaged grids by cryo-electron tomography (cryo-ET) and reconstructed tomograms using Etomo [33]. For both GS and hNuc, the lipid monolayer grids resulted in an equal or increased number of protein particles in the ice compared to the holey carbon grids (Figure 2A-B, top panels). For each tomogram, we calculated ice thickness and counted the number of particles in the field of view (Figure 2C). In general, the Ni-NTA lipid affinity grids had thinner ice as compared to holey carbon Quantifoil TEM grids (32+/-2 nm vs 51+/-14 nm for hNuc and 28+/-2 nm vs 42+/-3 nm for GS). By looking at an XZ slice through the tomograms (Figure 2A-B, bottom panels), it can be observed that Ni-NTA grids result in a single layer of protein. In contrast, holey carbon grids are seen to have particles at both AWIs, as seen before[3, 34].

**Figure 2.**
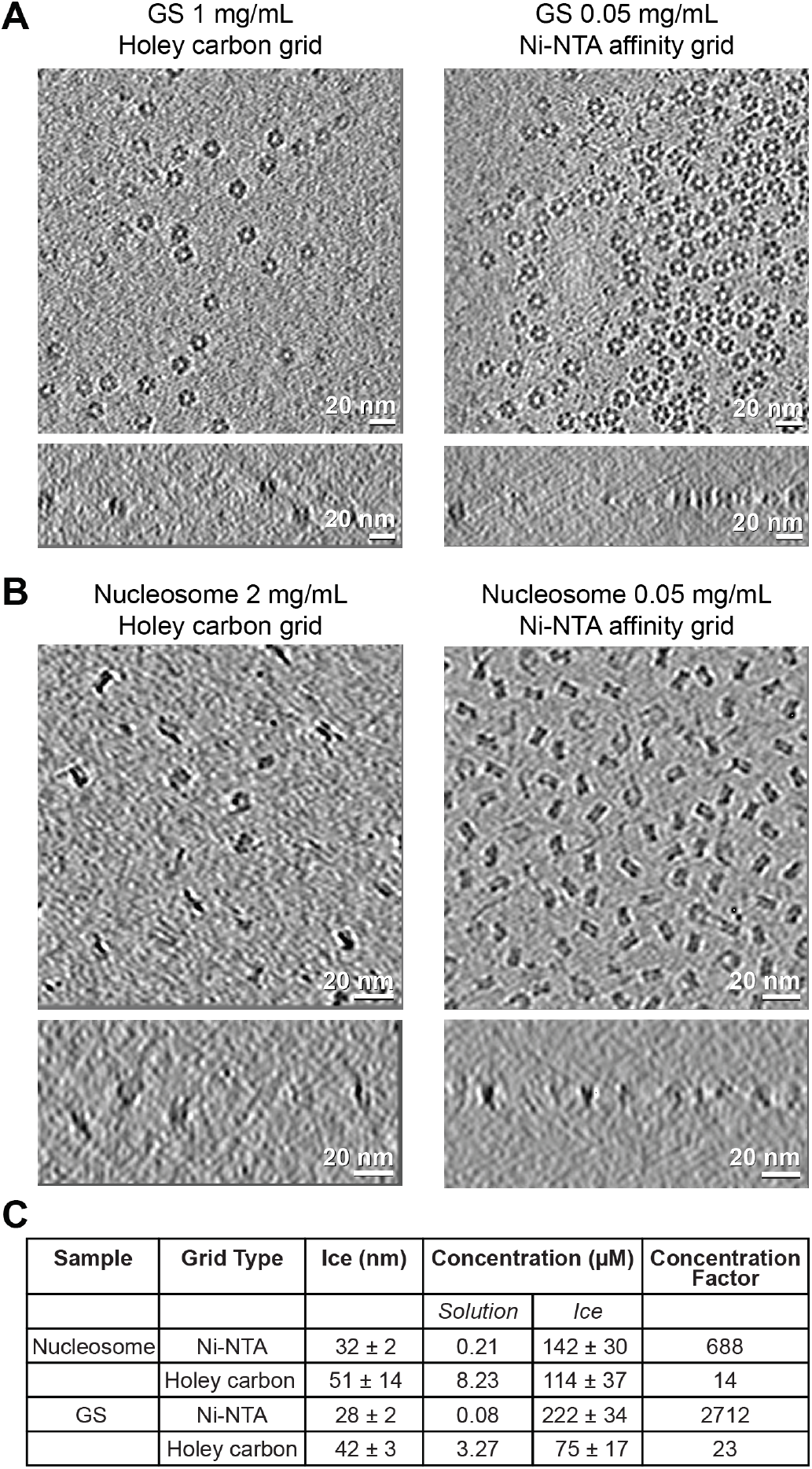
His-tagged protein complexes are concentrated on the lipid surface away from AWI in a layer of thin vitreous ice. Cryoelectron tomography of GS **(A)** and hNucleosomes **(B)** on Ni-NTA affinity grids. Average tomographic slices in XY or XZ, scale bar 20 nm. **(C)** Particle density, average ice thickness, and concentration factor for Ni-NTA and holey carbon grids. Concentration factor was calculated by comparing concentration in the tomogram compared to the initial concentration of the sample.

Finally, using ice thickness and particle number, we calculated the concentration of particles on the grid. By comparing the starting concentration of each sample in solution and the final on-grid concentration, we show that the Ni-NTA monolayer has a concentrating effect of 688-fold (hNuc) and 2712-fold (GS), compared to 14-fold and 23-fold, respectively, for holey carbon grids. This analysis shows that lipid monolayers can highly-concentrate samples away from the AWI and produce a uniform monolayer of particles embedded in thin ice.

### Ni-NTA lipid monolayers are amenable to high-resolution structural analysis

We next sought to determine if Ni-NTA monolayers are suitable for determining high-resolution single-particle reconstructions. To serve as a suitable grid support, the lipid monolayer must uniformly cover the grid to provide a sufficient number of micrographs, which each must have suitably thin ice to preserve high-resolution signal. Furthermore, it is unclear if the non-random distribution on the grid surface via the His-tag would lead to a preferred orientation. In the case of the dodecameric GS, we observe a 100% preferred orientation on the grid, presumably from the multiple His-tags all residing on the co-planar face of the ring-like structure (Supp. Fig. 2) We therefore chose our hNuc sample for full structural analysis as the well-known “side” and “disc” views were readily observable in our test data.

We collected a dataset consisting of 3,332 micrographs on a Talos Arctica 200-keV cryo-TEM equipped with a Gatan K3 direct electron detector. The dataset was processed in Relion to a final gold-standard FSC_0.143_ resolution of 2.74 Å with C1 symmetry and 2.59 Å with C2 symmetry (Figure 3A, Supp. Fig. 3, Table 1). High-resolution features consistent with this FSC value are observable, such as recognizable sidechain densities and fully resolved DNA bases (Figure 3B). As shown in the angular distribution of the C1 refinement (Figure 3C), there is a bias in the distribution of particles with respect to the monolayer, as the complex is tethered via a His-tag on the N-terminus of histone H2A. However, this analysis shows that the flexible nature of the tether still allows for a sufficiently random orientation distribution relative to the lipid monolayer (Figure 3D).

**Table 1.**
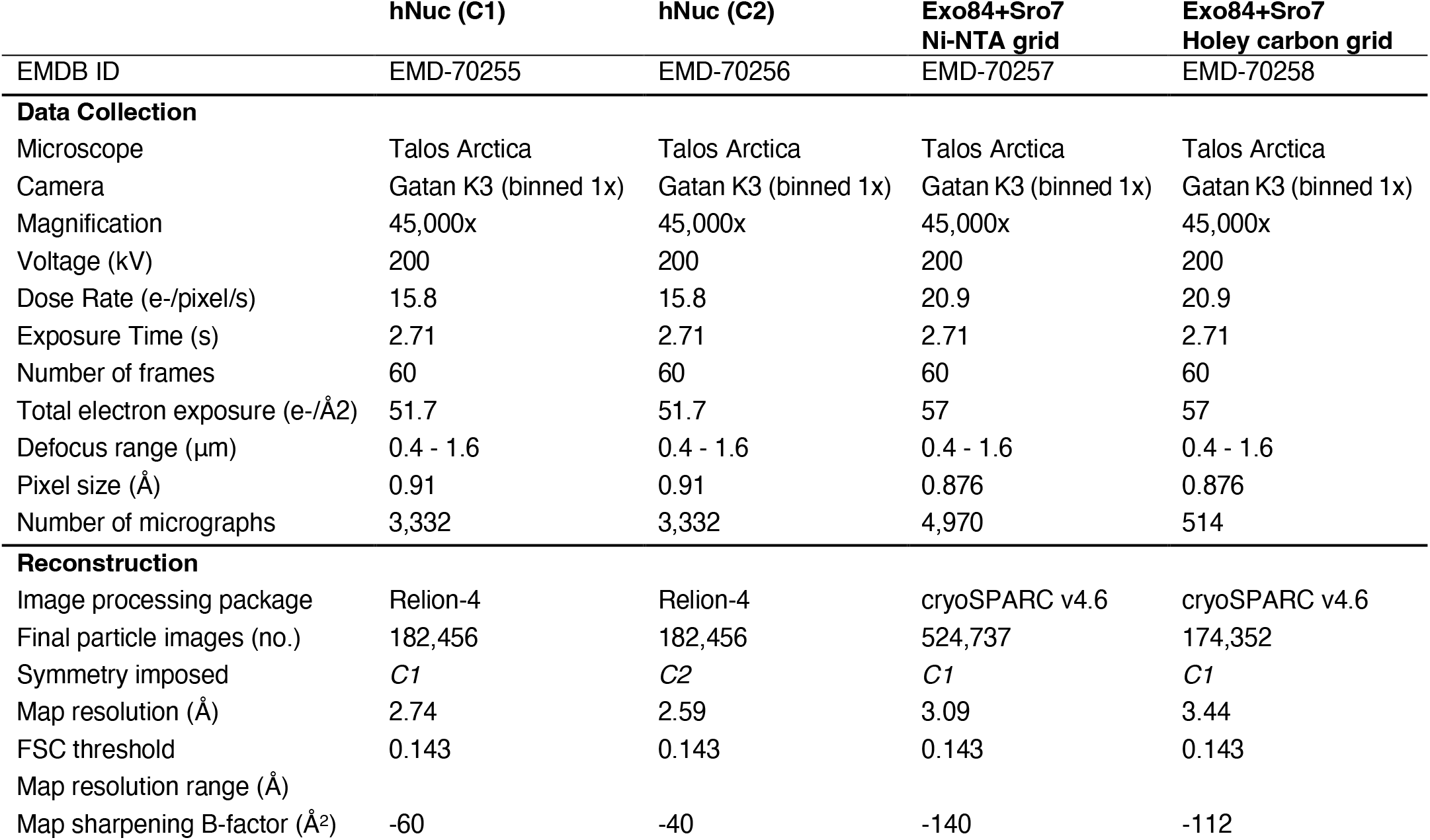
Cryo-EM map statistics

|  | hNuc (C1) | hNuc (C2) | Exo84+Sro7<br>Ni-NTA grid | Exo84+Sro7<br>Holey carbon grid |
| --- | --- | --- | --- | --- |
| EMDB ID | EMD-70255 | EMD-70256 | EMD-70257 | EMD-70258 |
| **Data Collection** |  |  |  |  |
| Microscope | Talos Arctica | Talos Arctica | Talos Arctica | Talos Arctica |
| Camera | Gatan K3 (binned 1x) | Gatan K3 (binned 1x) | Gatan K3 (binned 1x) | Gatan K3 (binned 1x) |
| Magnification | 45,000x | 45,000x | 45,000x | 45,000x |
| Voltage (kV) | 200 | 200 | 200 | 200 |
| Dose Rate (e-/pixel/s) | 15.8 | 15.8 | 20.9 | 20.9 |
| Exposure Time (s) | 2.71 | 2.71 | 2.71 | 2.71 |
| Number of frames | 60 | 60 | 60 | 60 |
| Total electron exposure (e-/Å<sup>2</sup>) | 51.7 | 51.7 | 57 | 57 |
| Defocus range (μm) | 0.4 - 1.6 | 0.4 - 1.6 | 0.4 - 1.6 | 0.4 - 1.6 |
| Pixel size (Å) | 0.91 | 0.91 | 0.876 | 0.876 |
| Number of micrographs | 3,332 | 3,332 | 4,970 | 514 |
| **Reconstruction** |  |  |  |  |
| Image processing package | Relion-4 | Relion-4 | cryoSPARC v4.6 | cryoSPARC v4.6 |
| Final particle images (no.) | 182,456 | 182,456 | 524,737 | 174,352 |
| Symmetry imposed | *C1* | *C2* | *C1* | *C1* |
| Map resolution (Å) | 2.74 | 2.59 | 3.09 | 3.44 |
| FSC threshold | 0.143 | 0.143 | 0.143 | 0.143 |
| Map resolution range (Å) |  |  |  |  |
| Map sharpening B-factor (Å<sup>2</sup>) | -60 | -40 | -140 | -112 |

**Figure 3.**
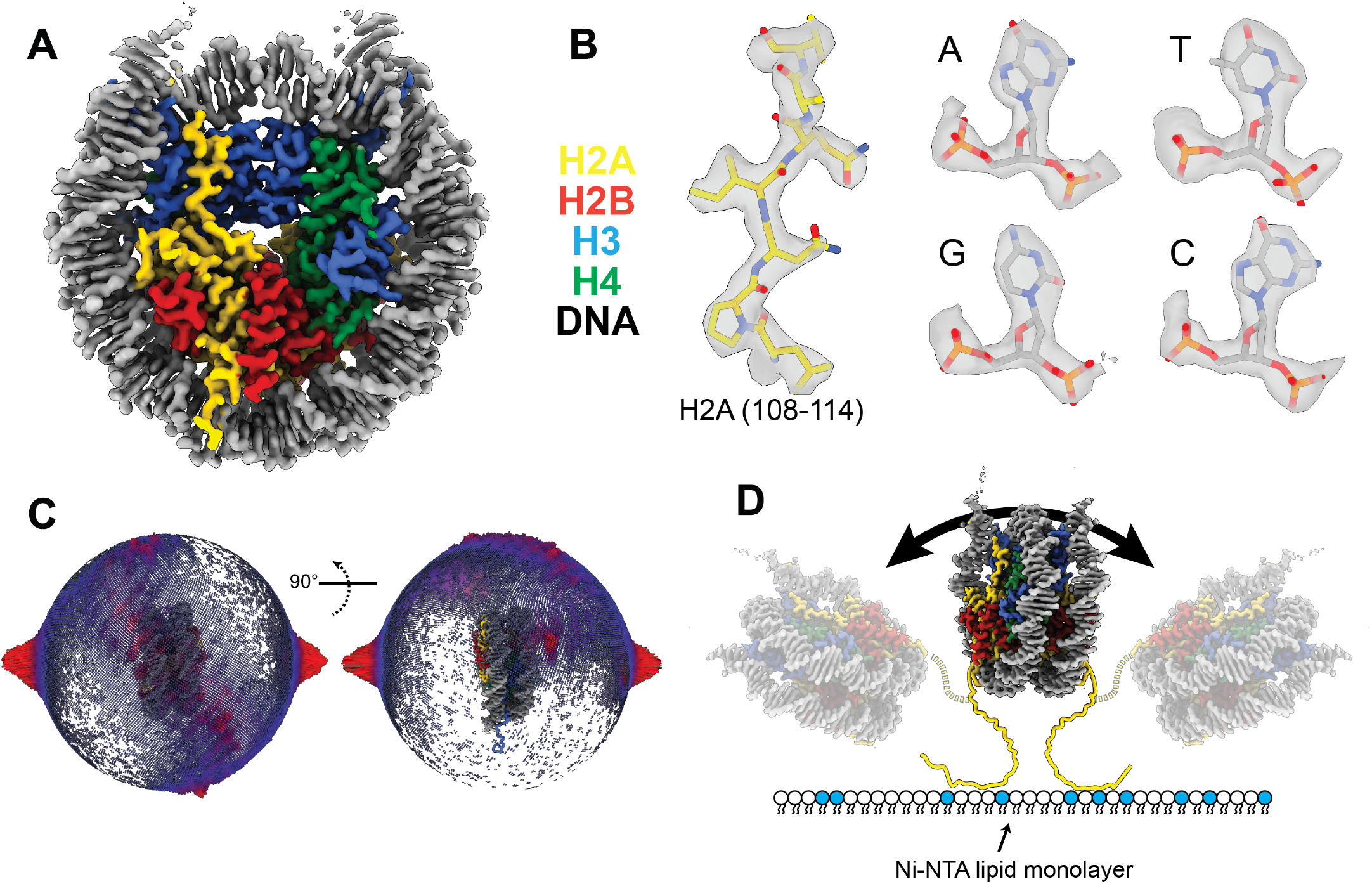
Single-particle cryo-EM of nucleosomes on Ni-NTA affinity grids. **(A)** cryo-EM density of the hNuc reconstruction with a final GSFSC of 2.6 Å. Coloring as labeled. **(B)** Cryo-EM density of high-resolution protein and DNA features in the hNuc map. **(C)** Angular distribution plot of a C1 reconstruction. **(D)** Schematic of hNuc tethered to a Ni-NTA monolayer, highlighting the flexibility of the tail to enable multiple views of the complex.

### On-grid complex assembly using lipid monolayers

A common problem when determining cryo-EM structures of protein complexes is that the final dataset contains a mixture of bound and unbound complexes. One strategy to overcome this issue would be to tether one binding partner to the Ni-NTA lipid monolayer and form complexes on grid. Unbound complex could be washed away, and particles still remaining on the grid would presumably be enriched for the fully formed complex. To test this on-grid binding strategy, we tested if one His-tagged protein could recruit an un-tagged binding partner to the grid surface (Figure 4A). We have previously determined structures of this complex, which comprises the clathrin adaptor complex AP-2 and the endocytic regulator SGIP [35]. To test the recruitment to grids, we pre-incubated grids with His-SGIP or a buffer control, then washed and incubated with 0.1 mg/mL AP-2. This is approximately 40-fold lower concentration compared to the conditions used for our previous structure using holey carbon grids. As shown in Figure 4B, AP-2 is robustly recruited to grids after pre-seeding with His-SGIP.

**Figure 4.**
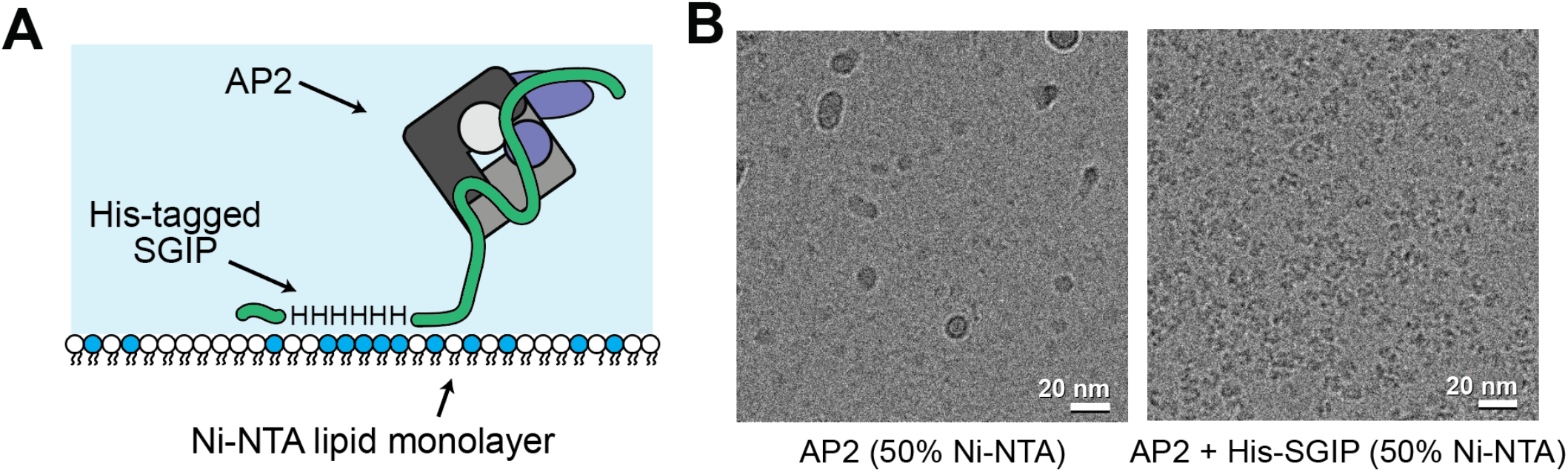
Affinity grids can be used to build complexes on-grid. **(A)** Schematic of the AP-2 clathrin adaptor binding to His-tagged SGIP on a Ni-NTA affinity grid. **(B)** Comparison of AP-2 recruitment to an Ni-NTA grid alone (left) or in the presence of His-tagged SGIP.

### High-resolution structure determination of a sub-100 kDa protein

To test the general nature of our protocol, we next turned to a protein complex of interest in our research group, a cocomplex between the Exocyst subunit Exo84 and the yeast tomosyn homolog, Sro7. The crystal structure of yeast Sro7 has been previously reported [36]. The binding of Sro7 to the N-terminus of Exo84 activates the tethering function of the Exocyst complex [37]. As our Exo84 construct contains an N-terminal His-tag, we prepared Ni-NTA affinity grids using pre-formed His-Exo84/Sro7 complex (Figure 5A). We collected 4,970 micrographs and processed the dataset in cryoSPARC [38] to a a final gold-standard FSC_0.143_ resolution of 3.1 Å (Figure 5B, Supp. Fig. 4, Table 1). Unfortunately, we do not observe density for Exo84, presumably because the interaction occurs within the unstructured N-terminal tail of Sro7. The resolved region of our map shows only the tandem WD40 domains of Sro7, with a molecular mass of ∼97 kDa. The secondary structure of Sro7 is nearly entirely beta sheet, and our structure shows well resolved beta strands (Figure 5C). Overall, these data demonstrate that this technique is amenable to proteins without symmetry, as well as proteins below 100 kDa.

**Figure 5.**
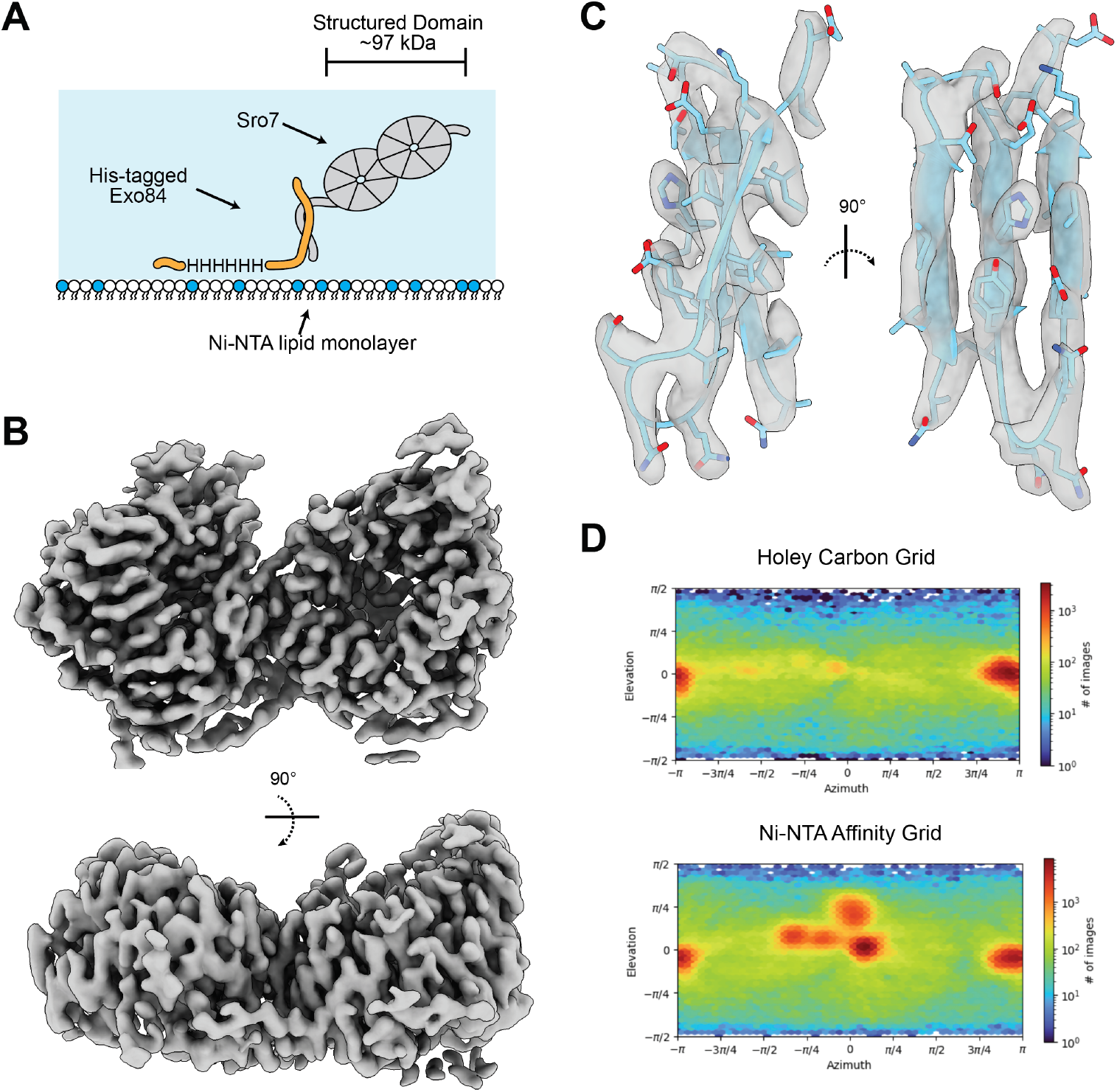
Single-particle cryo-EM of a sub-100 kDa protein using Ni-NTA affinity grids. **(A)** Schematic of the His-Exo84 + Sro7 sample used with the Ni-NTA affinity grid. **(B)** cryo-EM volume of the 3.1 Å reconstruction, showing the dual beta propellors of Sro7. **(C)** Zoomed-in view of a beta sheet, showing residues 90-116 of Sro7. Note the separation of the beta strands. **(D)** Angular distribution plots for the two grid types compared in this study.

In addition to our Ni-NTA grids, we also prepared holey carbon cryo-EM grids of the Exo84/Sro7 complex. In general, these grids were less reproducible and suffered from having either too few particles, or extremely dense distribution with particles at both AWIs. One test dataset of 514 micrographs reached a final gold-standard FSC_0.143_ resolution of 3.5 Å, with features consistent with this resolution (Supp. Fig. 5, Table 1). While both datasets have a predominant “side-view” and the final reconstructions show mild anisotropy, they have similar sphericity values (0.738 and 0.779) and directional FSC profiles (Supp. Figs. 4H and 5F). However, the angular distribution plots from the final reconstructions show a clear difference (Figure 5D), with several views being much more highly populated in the Ni-NTA dataset. While this did not meaningfully change the anisotropy of the final reconstruction, the use of a Ni-NTA support had a clear effect on the angular distribution, suggesting that this technique may be useful in overcoming preferred orientation problems in other cases.

## Discussion

We present here a simple and generalizable protocol for creating cryo-EM grids coated with a Ni-NTA lipid monolayer and benchmark the utility of this protocol with modern cryo-EM methods. Using multiple protein complexes, we demonstrate benefits of this technique as compared to holey carbon TEM grids, including (1) concentrating particles away from the AWI, (2) greater particle absorption to the grid (i.e. on-grid higher concentration factor) and (3) thinner ice (∼30 nm) suitable for high-resolution data collection. Overall, this is a simple and straightforward method with application to a variety of samples for high-resolution structural analysis.

### Practical considerations when using Ni-NTA Affinity grids

We have created a detailed protocol that is available in the supplementary materials. The protocol takes between 3-6 hours, depending on the number of grids prepared, and can be done in a general laboratory setting with equipment typically found in a cryo-EM laboratory or core facility [39]. The preparation of the Ni-NTA films requires a few hours of extra time to prepare prior to specimen vitrification. Lipid monolayers can be prepared in advance, dried, and stored in a dessicator for up to several months, as reported for 2D crystals of streptavidin monolayers [20]. We have had success in using pre-fabricated 20% Ni-NTA monolayer grids stored up to several weeks in a dry-keeper. For use, grids were simply rehydrated with buffer (150 nM NaCl and 20 mM HEPES pH 7.4) prior to sample incubation and cryo-plunging. In general, prefabrication of lipid affinity grids enables greater accessibility to this technique, with less time spent preparing lipid monolayers for each experiment, making the protocol more streamlined.

### Lipid monolayers and preferred orientation

Before our systematic analysis, it was unclear if the presence of the His-tag at one terminus of the protein would bias the orientation of the particle relative to the lipid monolayer. Indeed, Ni-NTA-containing membrane tubules have been used for the cryo-EM analysis of His-tagged proteins [40], with a primary benefit being the lack of orientation bias from using a cylindrical membrane. Of the three samples used in this study, only one had strong preferred orientation for monolayer Ni-NTA affinity grids. The GS adopted a 100% preferred orientation on Ni-NTA grids, although we also see a preferred orientation in open holes. We speculate that a combination of the D6 symmetry and orientation of the His-tags to the same plane led to preferred orientation on the lipid films, as this was not observed with hNuc or Sro7. Indeed, the hNuc sample has two tags, which do not lead to preferred orientation. The distribution, geometry, and location of the poly-His tags may therefore be critical to how the particle adheres to the lipid monolayer. While we did not systematically test linker length, the hNuc sample, which generated the highest resolution map in our examples, included a 16 amino-acid linker. Practically, this linker is longer, as histone tails are unstructured, giving our sample ∼26 amino acids of disordered sequence between the His-tag and the nucleosomes. Our successful hNuc reconstruction provides a benchmark for designing His-tagged proteins to use in our protocol.

### Ni-NTA monolayers can highly concentrate particles on the grid

One potential benefit of affinity grids is the ability to concentrate low abundance samples onto the grid. Using cryo-electron tomography (cryo-ET), we calculated concentration factors for two His-tagged protein complexes on Ni-NTA grids compared to holey carbon grids. We found that Ni-NTA lipid films concentrate proteins by a factor of 688 and 2712 for hNuc and GS, respectively. Cryo-EM studies, prior to the advent of direct-electron detectors, reported the isolation of dilute macromolecular complexes including ribosomes, aquaporin-9, and a co-complex of transferrin receptor bound to transferrin (Tf-TfR) from crude cell extracts using lipid monolayers [29, 41]. Based on the findings summarized in this study, we expect that on-grid purification could yield near-atomic resolution cryo-EM maps utilizing Ni-NTA affinity grids. The addition of low mM imidazole, or multiple wash steps in buffer containing high salt, or other additive would aid in removal of non-specific bound proteins to the lipid film.

### High resolution cryo-EM structures of protein complexes on Ni-NTA lipid monolayer

Using our affinity grid approach, we show that structural analysis to high resolution (2.6 Å in the case of hNuc) and for samples under 100-kDa (3.1 Å resolution for the 97-kDa dual WD40 motif of Sro7) is possible. For hNuc, the resolution of the map is comparable to published cryo-EM maps in the EMDB with a resolution range of 2.3 to 4.3[15, 42-44]. Importantly, nucleosome samples are often cross-linked to increase the stability of the complex. Our complex was prepared in the absence of chemical cross-linking, suggesting that lipid monolayers promote complex stability through added protection from the AWI, as reported for samples concentrated on graphene oxide [15].

An important consideration for high-resolution analysis is the thickness of the ice on the cryo-EM grid. In our analysis, the ice was thinner for the Ni-NTA affinity grids as compared to Quantifoil TEM grids, by a factor of ∼1.5. We reason that the thinner ice and distribution of particles as a single monolayer would aid in the acquisition of higher-resolution data for both single-particle analysis and cryo-ET subvolume averaging [45-48]. In the case of tilting to overcome a preferred orientation [49], tilting by 40° increases ice thickness by 1.3 times that relative to zero tilt. For samples with preferred orientation, if the Ni-NTA grid does not alleviate the preferred orientation, lipid monolayers may be of benefit to collect tilted data on the thinnest ice possible. However, at least for the case of Sro7, we see a change in the angular distribution on Ni-NTA monolayers as compared to open holes, suggesting that this method can be of benefit to certain samples in overcoming preferred orientation and obtaining isotropic reconstructions.

## Acknowledgements

We acknowledge Dr. Nathan Nicely of the UNC-CH Protein Expression core for purifying the GS used in this study. We acknowledge Mr. Jared Peck of the UNC-CH cryo-EM core facility for designing the forceps manifold. We acknowledge the UNC-CH cryo-EM core facility, which is in part supported by the National Cancer Institute of the National Institutes of Health under award number P30CA016086. The content is solely the responsibility of the authors and does not necessarily represent the official views of the National Institutes of Health. We acknowledge funding from the National Institutes of Health grant R35 GM150960 (RWB), R01 GM054712 (PB), R35 GM133498 (RKM) and Alfred P. Sloan Foundation grant G-2021-14197 (RWB).

## Supporting Materials

A full, step-by-step protocol for making lipid monolayer grids is included as a supplementary file.

The 3D printing plans for the forceps manifold is included as a supplementary file and also available at https://github.com/Rick-Baker/affinitygrids.

## Materials and Methods

### Protein expression and purification

*Staphylococcus aureus* glutamate synthase (*sa*GS) was expressed and purified according to published protocols [50] with a few modifications. Histagged GS expressed in *E. coli* C41(DE3) cells and purified by cobalt NTA column chromatography, followed by size exclusion chromatography using a Superdex 200 pg 16/60 column. Fractions enriched in GS (24.50 mg/ml) in buffer A (50 mM Tris (pH 7.5), 0.5 mM bME and 5% glycerol) were pooled and flash frozen in liquid nitrogen.

Human histones including Flag-His-tagged H2A (hH2A.D), H2B (hH2B.C), H3 (hH3.2), and H4 (hH4) were expressed, purified and assembled into nucleosomes essentially as described in [51]. Briefly, FLAG-His-hH2A.D/hH2B.C dimers were coexpressed in BL21(DE3)pLysS and purified using polyethylenimine (PEI) precipitation, while hH3.2 and hH4 were expressed individually, purified from inclusion bodies and refolded into hH3.2/hH4 tetramers. The 185 bp 601 nucleosome positioning DNA sequence was purified from E. coli as previously described [52]. Nucleosomes were reconstituted by gradient dialysis using H2A/H2B dimers, H3/H4 tetramers and 185 bp 601 DNA in the 2.8:1.0:1.0 ratio, respectively.

*Mus musculus* AP-2 and *Mus musculus* SGIP were purified as described [35]. Briefly, AP-2 was co-expressed on two plasmids in BL21 *E. coli* and purified with a GST tag on the alpha subunit. Our AP-2 construct includes alpha (1-621), beta-2 (1-591), mu-2 (FL), and sigma-2 (FL). SGIP residues 69-156 were purified with an N-terminal His tag after overexpression in BL21 *E. coli*.

*Saccharomyces cerevisiae* Sro7 and Exo84 (1-326) were purified essentially as described [37]. Briefly, full-length Sro7 was purified using the Tandem Affinity Purification (TAP)-tag system from yeast cultures overexpressing tagged Sro7 expressed behind *ADH1* promoter on a 2 micron plasmid. Histagged Exo84 (aa 1-326) was expressed in BL21 *E. coli* and purified using IMAC. Complexes of Sro7 and 6xHisExo84 were made up by mixing 0.8 µM Sro7 final concentration with 0.8 µM 6xHis-Exo84 final concentration in a binding buffer of 10 mM Tris pH 7.5, 150 mM NaCl, 2.5 mM MgCl_2_ and incubated for 1.5 hours at 25C. Complexes were put on ice and centrifuged at 13,000 x g for 10 minutes to remove any possible aggregates prior to placing on grids.

### Ni-NTA Affinity Grid preparation

Affinity grids containing Ni-NTA were prepared as previously described with a few modifications to the protocol[21, 25, 29]. Quantifoil R1.2/1.3 with gold mesh TEM grids (Quantifoil Micro Tools GmbH) were washed with chloroform and 95% ethanol and air dried over a snorkel before using. Lipids of either 18:1DGS-NTA (Ni Salt)1,2-dioleoyl-sn-glycero-3-[15] (nickel salt) or DOPC (18;1) 1,2-dioleoyl-*sn*-glycero-3-phosphocholine) were purchased from Avanti Polar Lipids (Alabaster, AL) and stored at -20 C°. Stock solutions of lipids at 1 mg/ml (w/v) in chloroform were allowed to incubate at room temperature for 1 hour before casting lipid mono-layers on reservoir buffer consisting of 50 or 10 mM HEPES (pH 7.5) and 0.15 M NaCl as described. In brief, 1 drop of castor oil was dropped onto the surface of reservoir buffer in a small (60 × 15 mm) Pyrex petri dish [20]. Next, 1 µl of Ni-NTA lipid cocktail was released onto the center of a castor oil droplet using a Hamilton syringe. The lipid monolayers were transferred to the carbon side of the Quantifoil grids and placed into an ice chilled humidity chamber. Two µl of specimen-specific buffer was added onto the lipid monolayers (carbon side of each grid) to prevent the films from drying out. Three µl of His-tagged protein samples were pipetted onto the lipid monolayers and allowed to incubate for at least 30 mins before plunge-freezing in liquid ethane:propane (60:40). Cryo-grids were prepared using a TFS Vitrobot Mark IV with the following settings: temperature 4 °C degrees, 95% humidity, blot force -10, blot time 4 seconds, wait time 0, drain time 0. Prior to plunge freezing each of the TEM grids was washed 3 times by touching the carbon side of the grid to 100-200 µl drops of specimen-specific buffer dispensed on parafilm. A detailed protocol with instructional images from the procedure is available as a supplementary file.

### Cryo-EM data collection

#### Single-particle cryo-EM

Single-particle cryo-EM data was collected with a Thermo Fisher Scientific (TFS) 200 keV Talos Arctica equipped with a AMETEK Gatan K3 DED as previously described [53]. Movies (TLZ compressed TIFF) were collected with SerialEM using the BIS method using a 5 × 5 multi shot array corresponding to maximum image shift of ∼ 7 µm [54]. The C2 lens was adjusted in diffraction mode under parallel illumination conditions using an evaporated aluminum TEM calibration grid (TedPella) [55]. Movies were collected at spot size 4, flux of 20 or 15.8 e_-_/unbinned pixels/second, and calibrated pixel size of 0.876Å or 0.91Å at the detector level. The 70 µm condenser lens aperture was inserted and the object lens aperture was retracted during data collection.

#### Cryo-ET

Tomography data was collected with a TFS 200 keV Talos Arctica equipped with a AMETEK Gatan K3 DED. Tilt series (maximum tilt range ± 60 with 3-degree increment) were collected with SerialEM using grouped dose-symmetric collection scheme, with 3 tilts group together until stage reached ± 45 degrees [56]. Total dose of each tilt series was at most 58 e_-_/Å_2_. Gained corrected movies were aligned with SerialEM using Alignframes, and processed with IMOD using fiducial less patch tracking method [33]. Tomographic reconstructions were generated using weighted back-projection algorithm and binned by a factor of 8 corresponding to a calibrated pixel size of either 1.4648 or 1.402 nm [57]. The CTF was estimated and corrected with IMOD CTFPlotter [58, 59]. The measured defocus ranged from 3.45 to 2.88 µm. Ice thickness and particle density were measured manually with the IMOD slicer window. Multiple tomograms were collected in the center of at least 3 grid squares for each sample used in the analysis.

### Cryo-EM processing

#### Nucleosome

Movies were imported into RELION 4.0 based on the beam image shifts (optics groups) used for data collection [60]. Motion correction and CTF estimation were conducted using the RELION implementation of MotionCor2 [61] and CTFFIND-4.1 [62], respectively, and only micrographs with an estimated resolution below 5 Å were used for further processing. Selected micrographs underwent RELION reference-free 2D classification using gradient-driven algorithm (3 rounds) to generate *de novo* 3D model of a nucleosome, which was then used in 3D classification. The best two 3D classes were combined for 3D refinement, followed by the per-particle CTF refinement. The refined model was subjected to another round of 3D classification, this time without alignment, to select highest-resolution particles. Selected particles underwent 3D-refinement, Bayesian polishing [63], per-particle CTF refinement and final 3D refinement using solvent-flattened FSCs. To further improve resolution, the final set of particles were extracted with a larger box size and refined with C2 symmetry following steps described above and in the Supp. Fig. 3. The final C1 reconstruction yielded a map with a nominal GSFSC resolution of 2.74 Å. The final C2 reconstruction yielded a map with a nominal GSFSC resolution of 2.59 Å. Map sharpening was performed using deepEMhancer [64]. 3D-FSC curves and sphericity values were calculated using the server at www.3dfsc.salk.edu.

#### Sro7-Exo84

All processing was performed in cryoSPARC v4.6.0 with similar processing schemes[38]. Movies were imported, motion corrected with dose weighting enabled, followed by patch CTF estimation and micrograph curation to remove outliers for CTF fit (worse than 4 Å) and defocus (outside of 0.5-2.0 µm). This resulted in a 514 micrograph dataset for open hole grids and 2,706 micrograph dataset for Ni-NTA grids. For each dataset, particles were picked with blob picker, followed by template picker using class averages from the initial round of 2D classification. Each particle dataset was extensively 2D classified, then combined and duplicated particles removed. This yielded a “clean” dataset of 174,452 particles for the open hole grid and 524,737 particles for the Ni-NTA grid. Particles were then sorted using *ab initio* model generation asking for 3 classes, which yielded a single class with clear structure resembling Sro7. The resultant particles were refined using non-uniform refinement [65], followed by 3D classification without alignment asking for 6 classes. Each class was refined independently, and the high-resolution classes were combined for a final refinement. This yielded a final reconstruction of the open-hole dataset with a gold-standard FSC_0.143_ resolution of 3.46 Å and a final reconstruction of the Ni-NTA dataset with a final gold-standard FSC_0.143_ resolution of 3.09 Å. Map sharpening was performed using deepEMhancer [64]. 3D-FSC curves and sphericity values were calculated using the server at www.3dfsc.salk.edu.

**Supplementary Figure 1.**
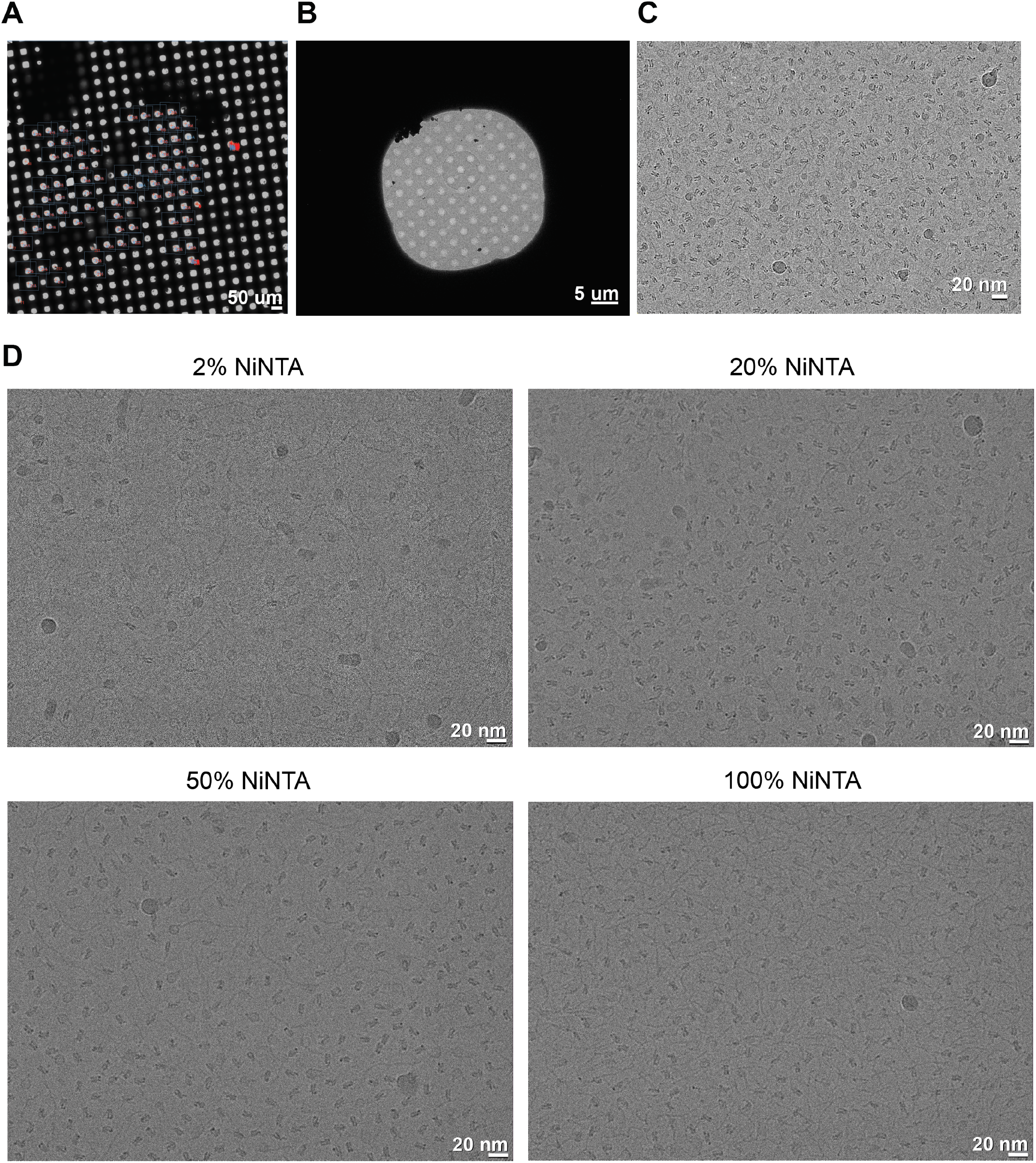
Example micrographs of Ni-NTA lipid grids and their effect on particle adsorption. **(A)** Atlas of the 20% Ni-NTA lipid affinity grid used for high-resolution data collection. Areas collected are indicated in blue squares. Scale bar 50 microns. **(B)** Micrograph of an example grid square collected from **(A)**. Scale bar 5 microns. **(C)** Dose-weighted, motion corrected micrograph collected on corresponding grid square, scale bar 20 nm. **(D)** Micrographs of different Ni-NTA affinity grids with various lipid compositions. Scalebar 20 nm.

**Supplementary Figure 2.**
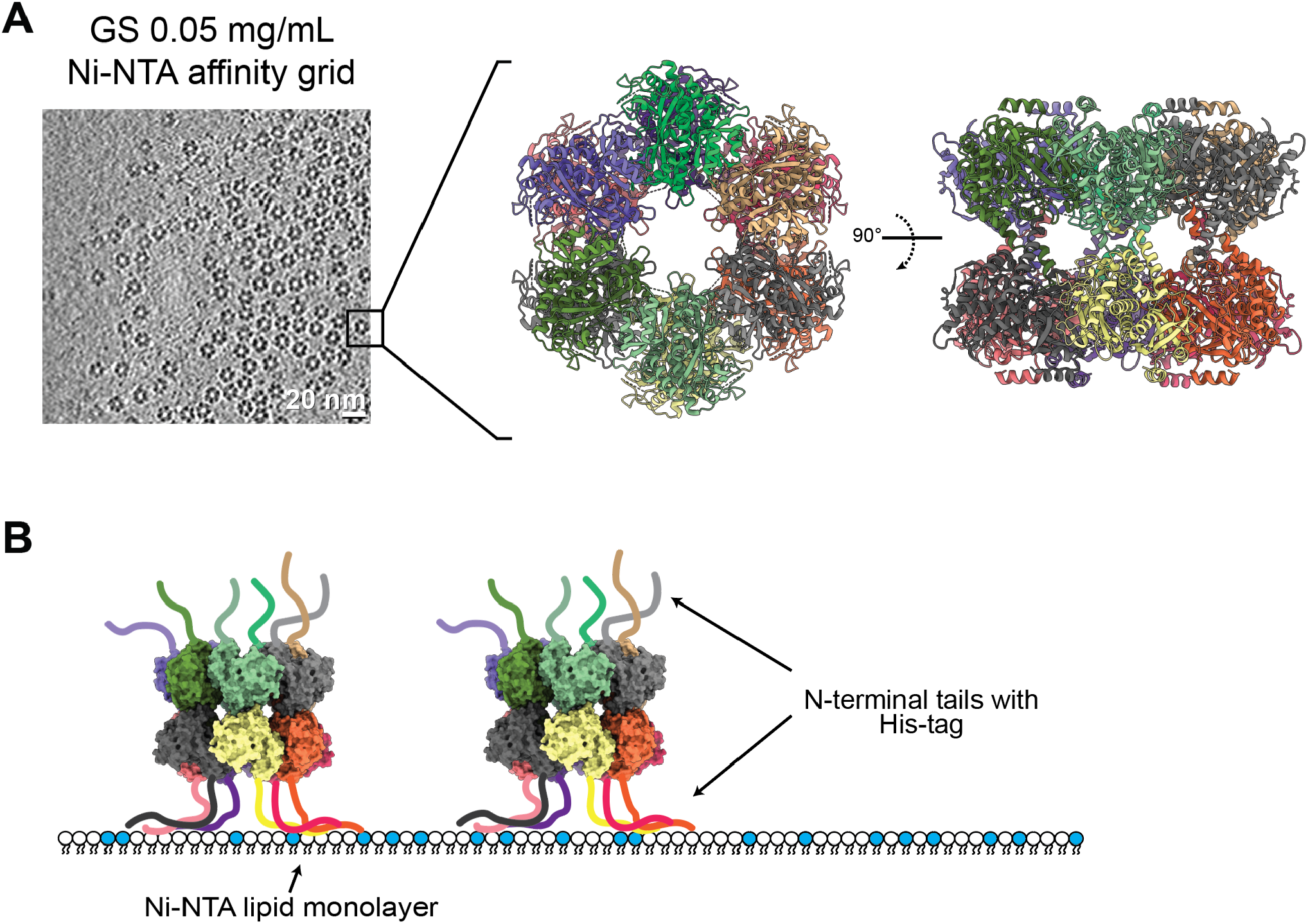
GS shows a preferred orientation on Ni-NTA affinity grids. **(A)** Example tomographic slice (scale bar 20 nm) of His-GS on Ni-NTA affinity grids, with a PDB model shown to highlight the hexameric ring structure of the particle. **(B)** Schematic of the double-ring structure of GS inducing a preferred orientation on a Ni-NTA lipid monolayer.

**Supplementary Figure 3.**
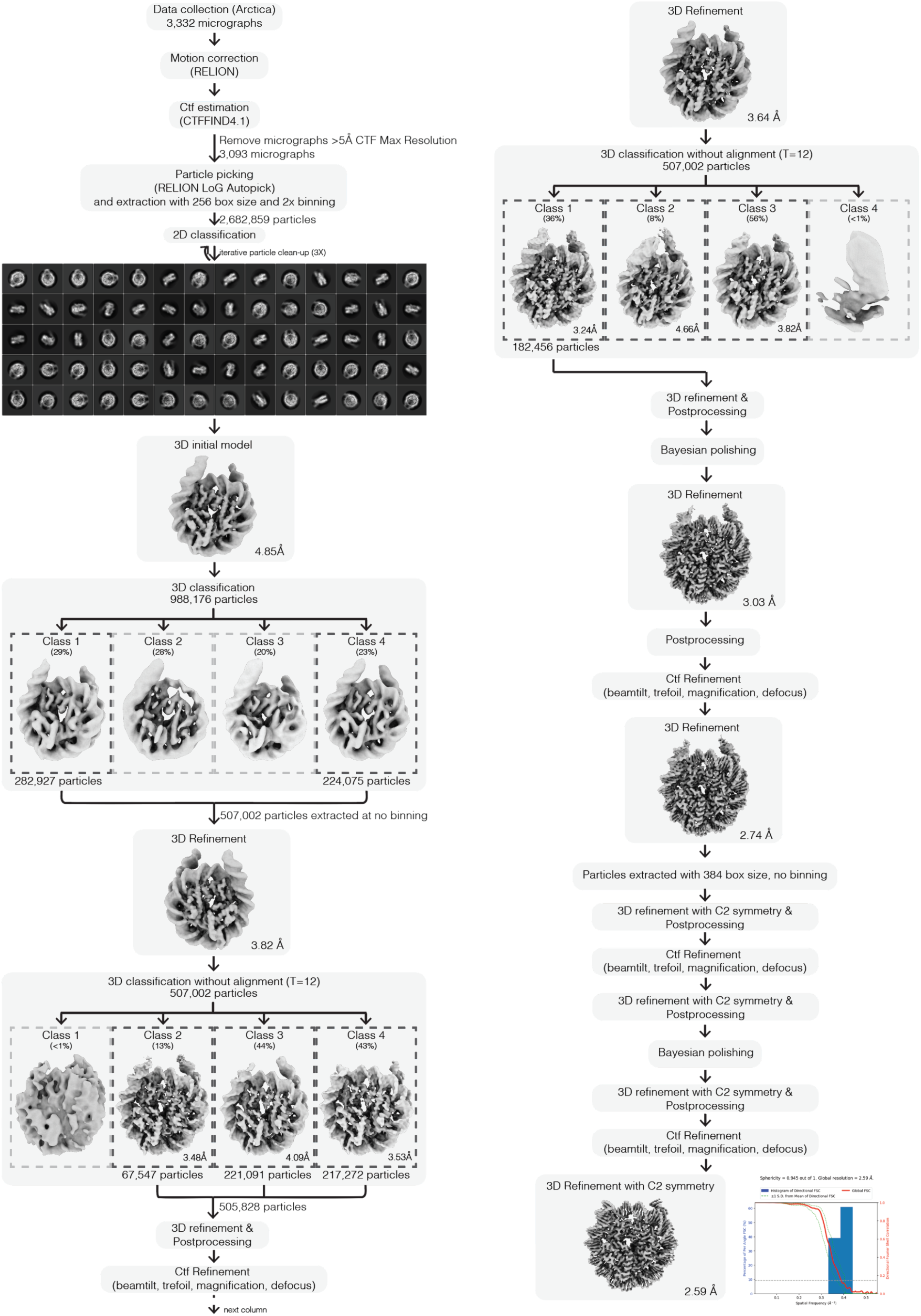
Cryo-EM processing pipeline for the human nucleosome complex on a Ni-NTA affinity grid. The hNuc cryo-EM dataset was processed end-to-end in Relion 4. Preprocessing, particle picking, 2D classification, 3D classification, 3D refinement, and particle polishing are highlighted in the schematic. Directional FSC curve and sphericity value is provided for the final C2 refinement, calculated using the www.3dfsc.slk.edu server.

**Supplementary Figure 4.**
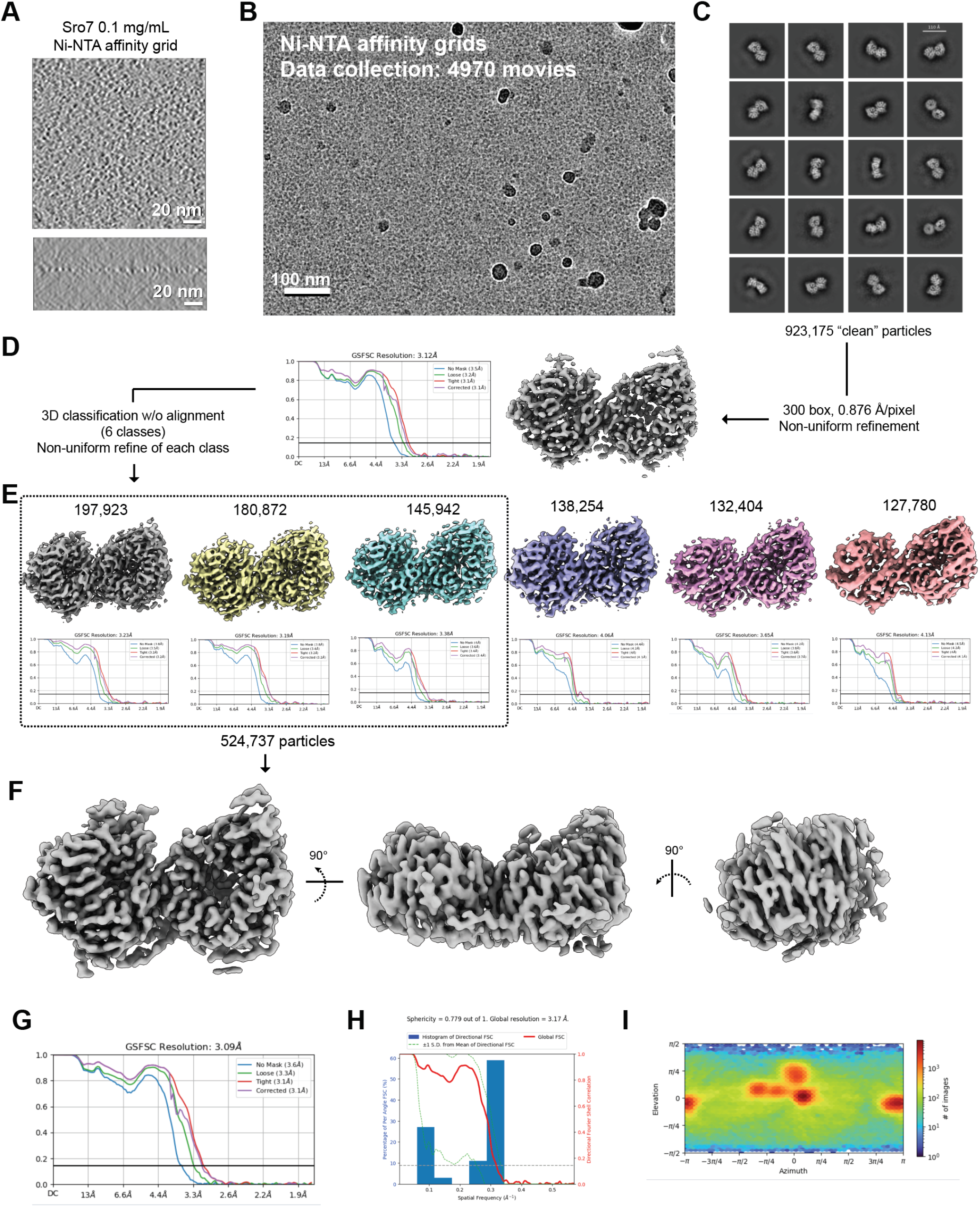
Cryo-EM processing pipeline for the His-Exo84/Sro7 complex on a Ni-NTA affinity grid. **(A)** Representative tomographic slices (XY and XZ) are shown for the Sro7 grid. **(B)** Example micrograph. Scale bar 100 nm. **(C)** 2D class averages from a later stage of classification. Multiple rounds of classification are used to clean particles selected with a variety of particle picking software. **(D)** ∼1m particles were refined then classified in 3D without alignment to reveal highly resolving sets of particles. **(E)** Each 3D class was subjected to Non-uniform refinement, and three high-resolution classes were combined for a finale round of Non-uniform refinement in **(F). (G)** FSC curves for the final 3D refinement. **(H)** Directional FSC curve and sphericity value, calculated using the www.3dfsc.slk.edu server. **(I)** Angular distribution plot from the final refinement. All processing was performed in cryoSPARC v4.6.

**Supplementary Figure 5.**
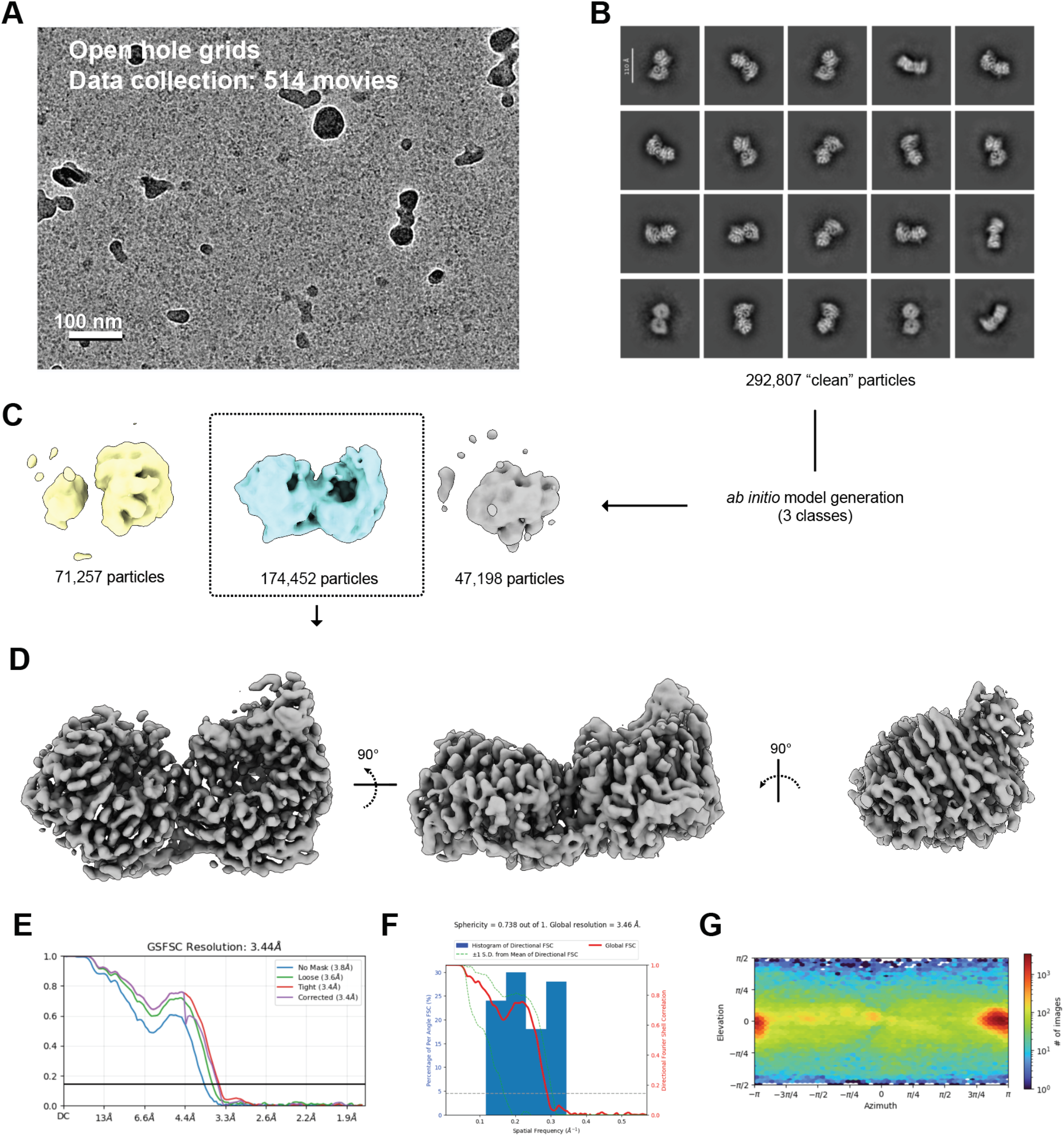
Cryo-EM processing pipeline for the His-Exo84/Sro7 complex on holey carbon grids. **(A)** Example micrograph. Scale bar 100 nm. **(B)** 2D class averages from a later stage of classification. Multiple rounds of classification are used to clean particles selected with a variety of particle picking software. **(C)** Particles were sorted using ab initio model generation with 3 classed. **(D)** Final refinement shown in 3 views. The map was sharpene used deepEMhancer. **(E)** FSC curves for the final 3D refinement. **(F)** Directional FSC curve and sphericity value, calculated using the www.3dfsc.slk.edu server. **(G)** Angular distribution plot from the final refinement. All processing was performed in cryoSPARC v4.6.

