## Supplemental Methods for "Nickel-NTA lipid-monolayer affinity grids allow for high-resolution structure determination by cryo-EM"

### Ni-NTA Affinity Grid Protocol

**Reagents and equipment:**

His-tagged protein stock.
Protein dilution buffer (sample specific).
HEPES buffer 10 mM (pH 7.5), 150 mM NaCl (buffer reservoir)

***Lipids***
18:1 DGS-NTA(Ni) (1,2-dioleoyl-sn-glycero-3-[(N-succinyl] (nickel salt)), supplied as 1 mg powder in sealed ampule (Avanti polar lipids, Cat. No. 790404).

DOPC (18;1) (1,2-dioleoyl-*sn*-glycero-3-phosphocholine), supplied at 2 mg/ml dissolved in chloroform in a a sealed ampule (Avanti Polar Lipids, Cat. No. 850375C-25mg).

***TEM grids***
Quantifoil Au 300 or 400 mesh R1.2/1.3 TEM grids.

***Humidity chamber***
The humidity chamber is a glass or pyrex baking dish (7 x 11in dish that is 3 in deep). The forceps holder is 3D printed (cat file included as a supplementary file, or available at https://github.com/Rick-Baker/affinitygrids).

***Syringes***
We use gas-tight Hamilton syringes in 2.5 µL, 5 µL,10 µL, 25 µL sizes for dispensing lipids (Hamilton, Cat. No CAL7632 with needle 7803-01, CAL87930, CAL80330, and CAL80430). The 2.5 µl syring was used to dispense the lipid mixture into castor oil for casting thin films. The needle was point style 4, angle 45. The other syringes were used to prepare lipid cocktails and were equiped with point style 2 needles.

**Procedure:**

**Bring the following reagents and equipment to the chemical hood.**

a. Chloroform ACS (Fisher Scientific, Cat. No. C298-500) and waste bottle.
b. Ethanol 99.5% anhydrous (Thermo Scientific, Cat. No. 61510-0010) or Ethyl Alcohol (95% ethyl Alcohol 5% isopropanol, Terrace Packaging) and a waste bottle.
c. Disposable pasteur pipettes 5 ¾ in. (Fisherbrand, Cat. No. 13-678-6A) and bulbs.
d. Volumetric flasks Glass A ASTM E288 (VWR) 25 and 1 ml.
e. Disposable culture tubes 6 x 50 mm Lime Glass (Fisherbrand, Cat. No. 14-958-A) and rack.
f. TEM grids in (Pyrex, 150 X 20 mm) petri dish containing Whatman #1 filter paper (GE Healthcare Life Sciences, Cat. No. 1001-090).
g. Anticapillary tip tweezers for handling Affinity grids (Dumont, Cat. No. 0203-N5AC-PO)
h. Fine tip forceps for TEM grids (Dummont, Cat. No. 0103-5-PS)
i. Two small (60 X 15 mm) Pyrex petri dishes.
j. DOPC and Ni-NTA lipids.
k. Parafilm (Parafilm M Laboratory sealing film) to cover glass culture tubes lipid cocktails.
l. Gas-tight 25, 10, 5, and 2.5 µL Hamilton syringes.

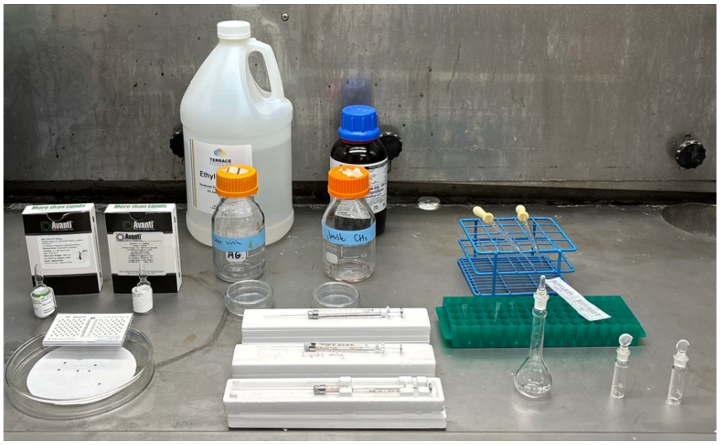

*Figure 1 - assembled components for grid making*

### Ni-NTA Affinity Grid Protocol

**Assemble humidity chamber.**

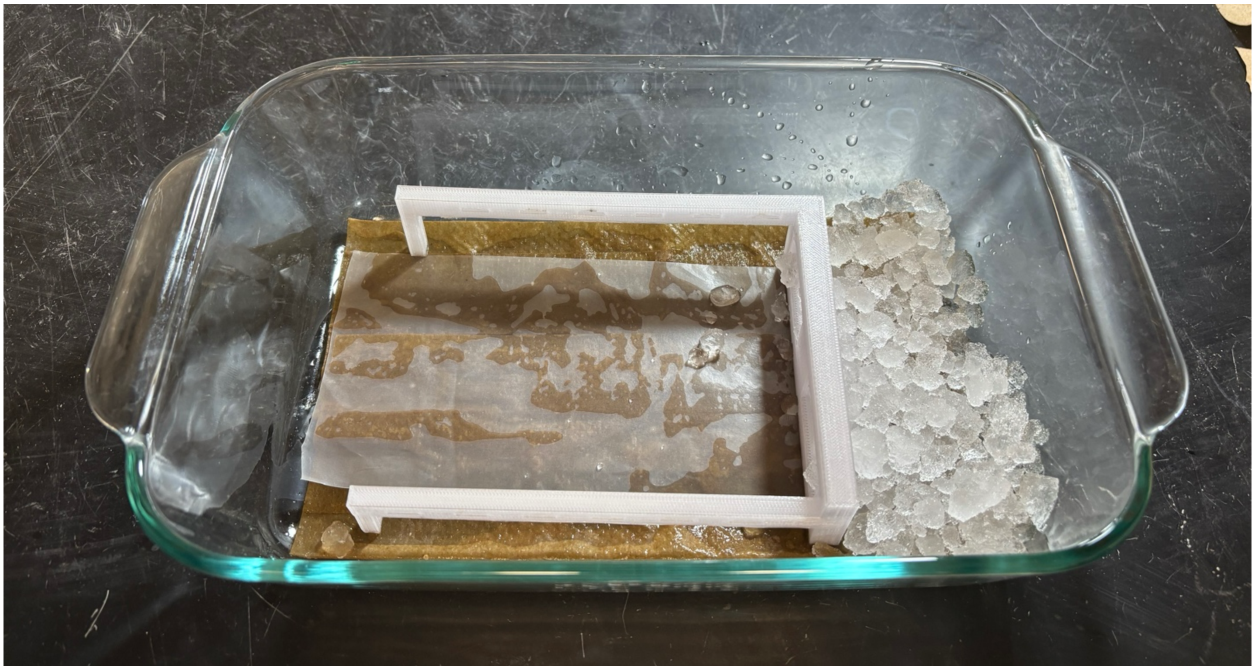

*Figure 2 - Assembled humidity chamber*

Place paper towels in Pryex baking dish. Soak with water, and cover with a pice of parafilm. Add ice to the side of the dish and place the forcep manifold inside. Cover with foil and let the chamber incubate to high-humidity. We normally setup the humidity chamber after preparing the lipid cocktails.

**Clean TEM grids in the chemical hood.**

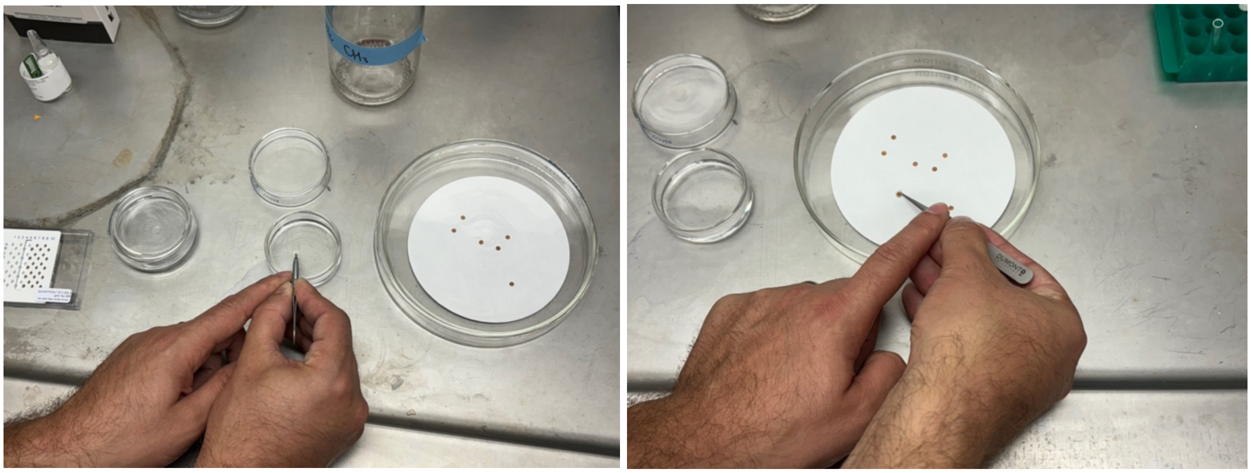

*Figure 3 - washing grids in organic solvent*

Grids need to be cleaned using organic solvents. First, wash two petri dishes, one with chloroform and one with alcohol (reagant alcohol or or 200 proof ethanol both work). Fill petri dishes with their respective solvents and use them to clean grids by dipping (vertically) first in chloroform, then into alcohol. Air dry grids by placing on Whatman filter paper with the carbon side facing up.

### Ni-NTA Affinity Grid Protocol

**Clean glassware and Hamiliton syringes with chloroform in the chemical hood.**
Wash Pasteur pipettes, Hamilton syringes, and glass culture tubes with chloroform. For pipettes and syringes, this requires pipetting and decanting several volumes of chloroform into a waste container. For tubes, pipette a few mLs of chloroform, swirl around, and pour the chloroform into a waste container. Air dry glassware and syringes in chemical hood.

**Preparing lipid stocks each 1 mg/ml (w/v).**
Dissolve 1 mg of Ni-NTA lipid in 1 ml of chloroform making a 1 mg/ml stock.  Add a small amount (less than 1 ml) of chloroform to the vial, mix then transfer into a 1 ml graduated flask. Fill graduated flask to the 1 ml mark with chloroform. Dilute the 25 mg/ml DOPC stock with chloroform to 1 mg/ml by taking 1 ml of stock and adjusting the final volume to 25 ml. Lipids can be stored in glass vials for several months in the -20°C freezer. We recommend adding dry nitrogen or argon gas over the tubes and sealing the tops with parafilm to keep out any air. Glass vials should have teflon gaskets or phenolic caps, as some gaskets/seals are not compatible with chloroform (e.g. Sigma cat. no. 27000 or FisherScientific cat. no. 03-338A).

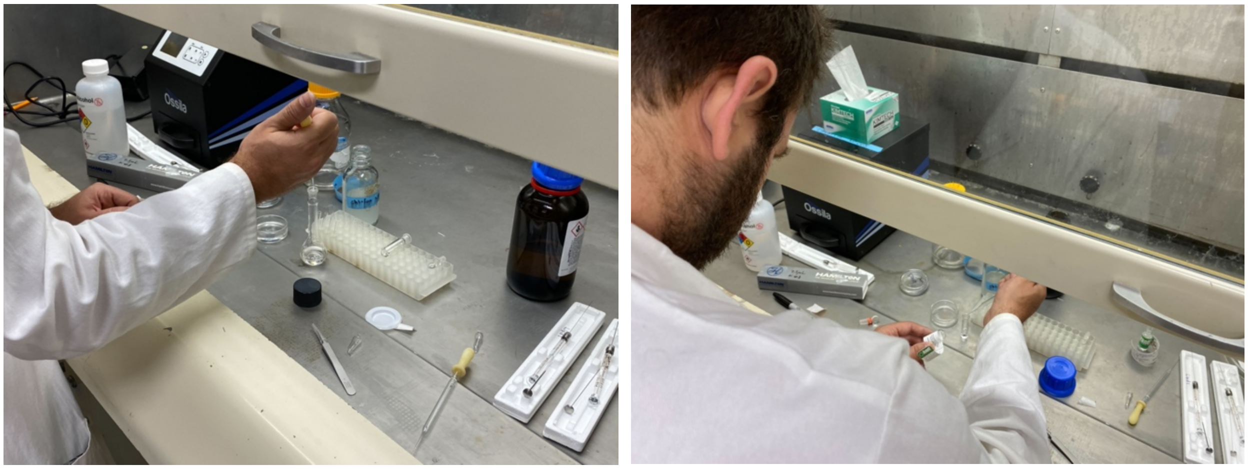


*Figure 4 - preparing lipid stocks*

**Prepare lipid cocktails in glass culture tube.**
Transfer ~ 50-100 µl DOPC and Ni-NTA to two glass culture tubes using the Hamilton syringe. Use prepared aliquots to make the lipid cocktail to avoid cross-contamating the lipid stocks. We recommend labeling the syringes "DOPC only" and "Ni-NTA only" as well. To make 20% (v/v) Ni-NTA cocktail add 40 µl of DOPC and 10 µl of Ni-NTA to a glass culture tube.  Mix gently with syringe. After making the lipid cocktail cover the top of the culture tubes with parafilm to avoid contamination and let equilibrate for about an hour in room temperature.

***Casting the lipid films and preparing the affinity TEM grids***

1. Before casting the lipid monolayer pickup each of the TEM grids with the anticapillary forceps (Dumont, Cat. No. 0203-N5AC-PO), this will save time.  Have the carbon side facing down and the bent tips of the forceps up, as shown in the image. Make sure to wash the forceps by dipping them in solvent and air dry before and after the experiment.

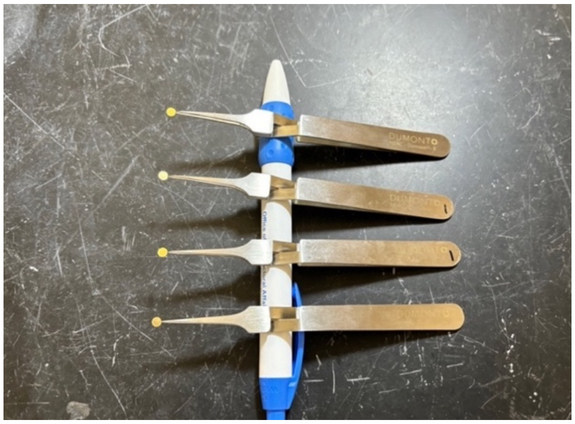

*Figure 5 - correct orientation for grabbing grids with forceps*

### Ni-NTA Affinity Grid Protocol

2. Cast lipid monolayer in small petri dish filled with HEPES buffer: add ~1 drop of castor oil to the dish, as shown in this image.

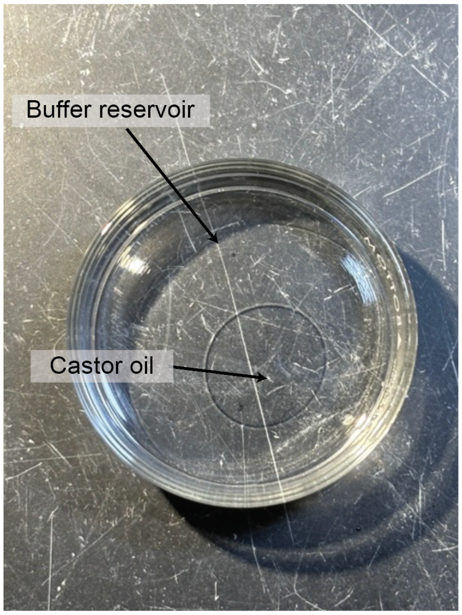


*Figure 6 - creating a castor oil droplet on buffer*

3. Orient the Hamilton syringe with the bevel facing down towards the petri dish. Dispense 1 μL of lipid, it should look like the image below (Figure 7).

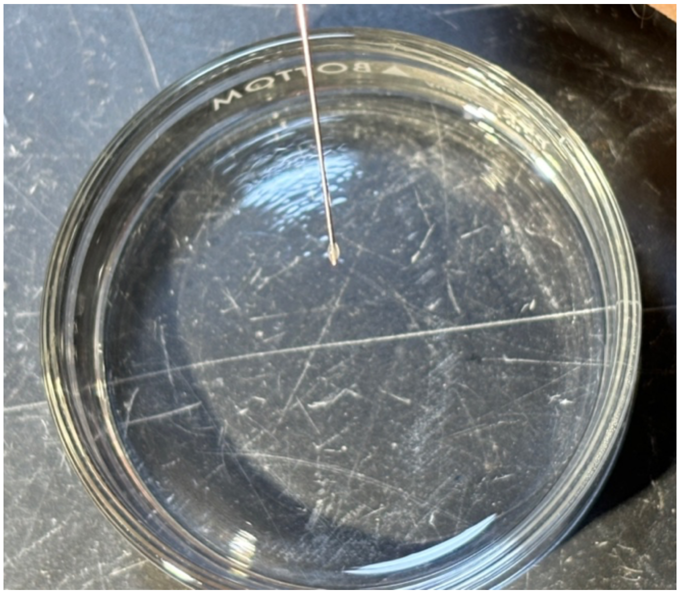

*Figure 7 - lipid droplet from syringe*

### Ni-NTA Affinity Grid Protocol

Carefully touch the lipid droplet to the surface of the center of the castor oil (Sky Organics). The lipid will be in the center of the dish as show in the image below (Figure 8). We recommend placing the petri dish over a blacktop table and illuminating the bench with a LED light.

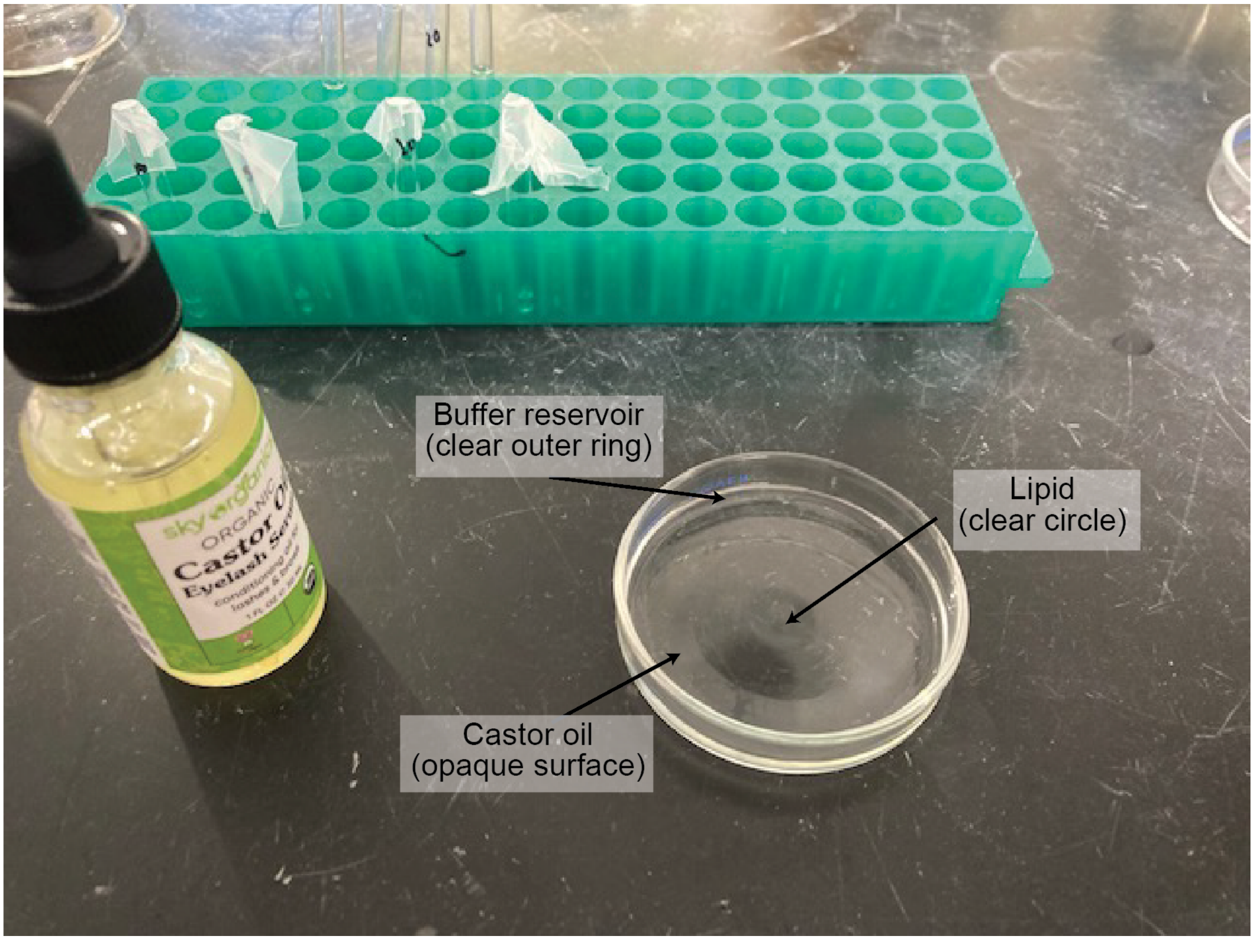


*Figure 8 - lipid monolayer on the castor oil droplet*

### Ni-NTA Affinity Grid Protocol

4. Transfer the lipid to the carbon side of a TEM grid using anticapiliary forceps as shown in this image.

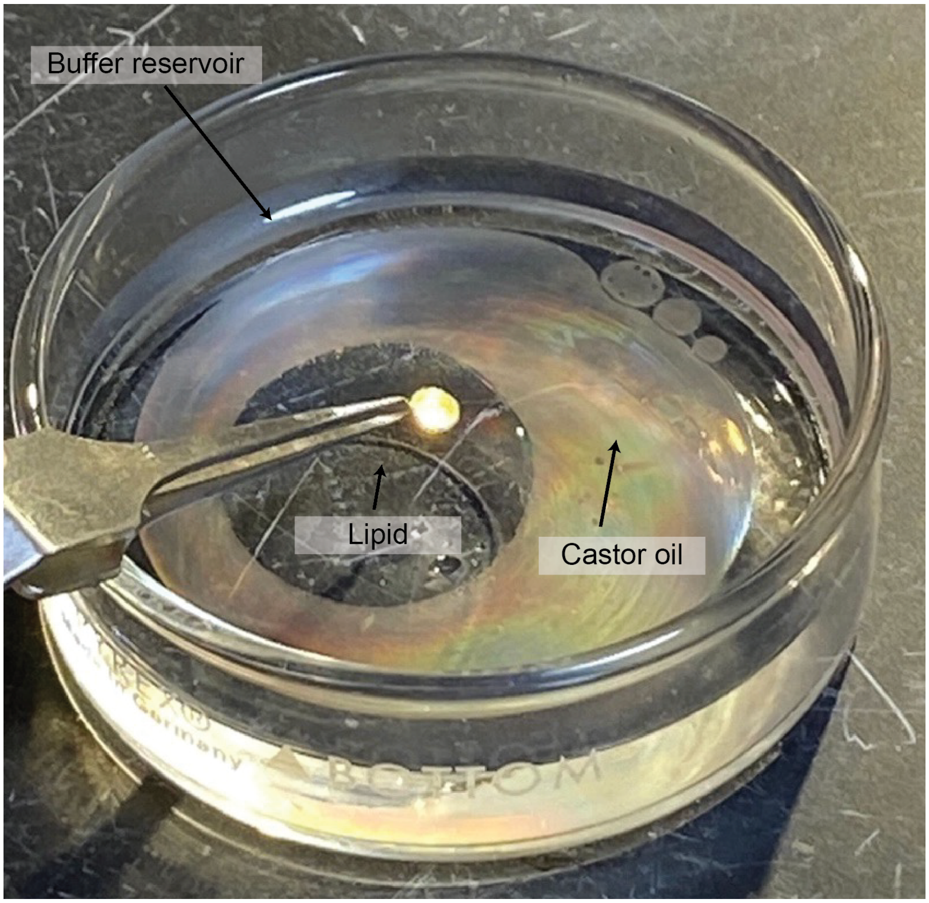


*Figure 9 - picking up a lipid monolayer with the carbon side of the grid*

The grid should have an intact lipid monolayer with buffer over it.

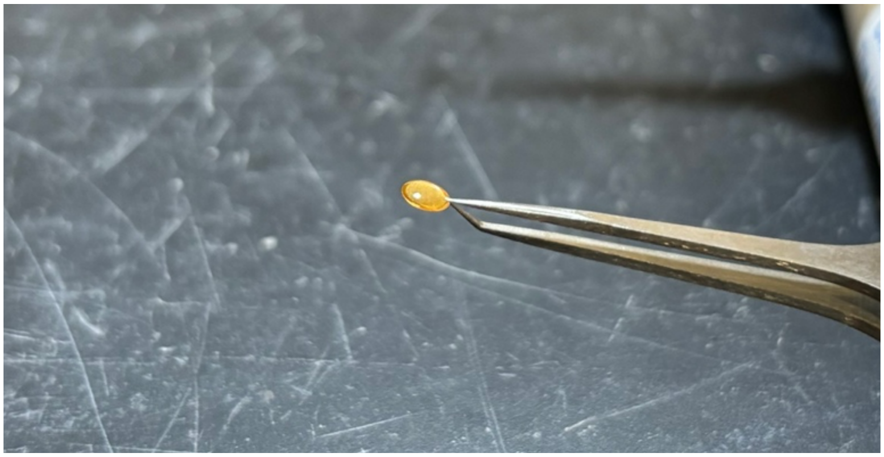

*Figure 10 - grid after lifting the lipid monolayer off the castor oil*

If you transfer the castor oil the liquid will ball-up on the surface of the TEM grid, as show in this image. If that happens do not use the grid, discard it.

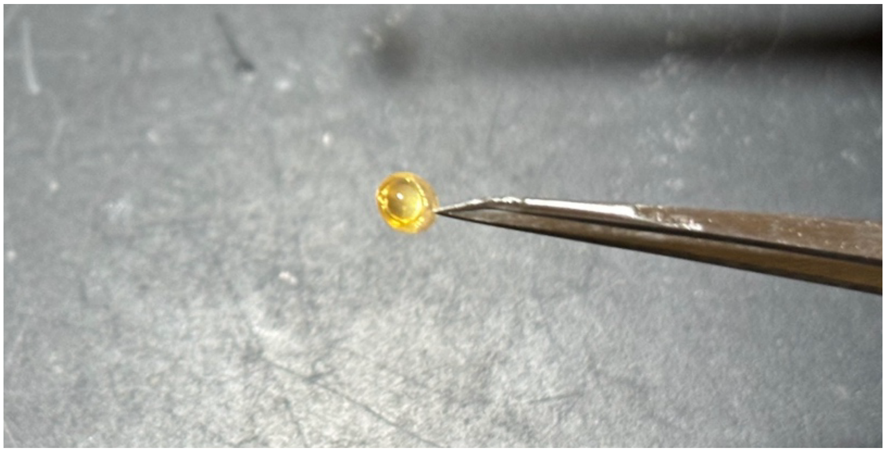

*Figure 11 - grid that has picked up castor oil, not lipid*

*Lipid affinity grids can be prepared in advance, dried, and re-hydrated with HEPES buffer. We have had success storing pre-fabricated lipid affinity grids for up to 2 weeks in a dry-keeper.*

5. Transfer forceps to the humidity chamber on the lab bench, filled with ice water and covered with paper-towels. Cover with lid during experiment but do not seal closed or use aluminum foil to cover to top. If you do not 3D print a forceps holder, any solid object that keeps the forceps off the ice will work, like a tube rack.

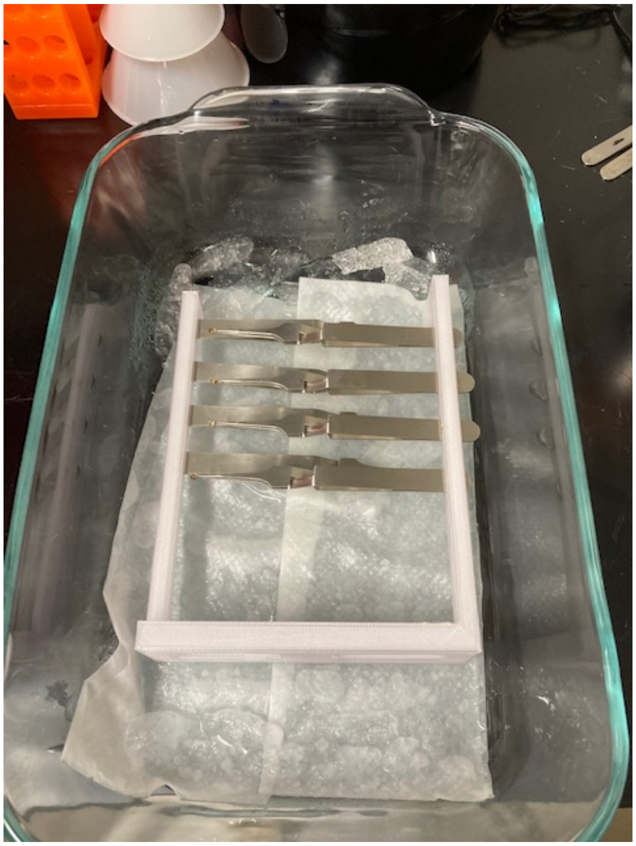

*Figure 12 - lipid-coated grids in the humidity chamber. There are 8 forceps in this image.*

### Ni-NTA Affinity Grid Protocol

6. With a pipette add 2 µl of protein dilution buffer onto the grids so lipid film does not dry out.

7. Dilute His-tagged protein to a final concentration of 0.05 – 0.1 mg/ml.

8. With a pipette add 3 µl of diluted His-tagged protein on each of the TEM grids.

9. Let forceps and grids incubate in high humidity for at least 30 mins.

**Cryo-plunging affinity grids with a Vitrobot**

*At this point the Vitrobot should be ready to use. We recommend setting up the Vitrobot during or before casting the lipid films. Make sure the humidity chamber and Vitrobot filter paper have incubated at high humidity for at least 30 mins. Also, cool the Vitrobot cryostat with liquid nitrogen and fill the ethane cup with ethane or an ethane:propane mix (40:60 %) and allow it to cool to ~190 °C (we use ethane:propane in the UNC cryoEM core). Have everything you need ready to cryo-plunge the affinity grids. Below are the conditions that work best on our Vitrobot Mark IV robot:*

a. Temperature 4 °C
b. 95% Humidity
c. Blot force -10
d. Blot time 4 seconds
e. Wait time 0 seconds
f. Drain time 0 seconds
g. Skip apply sample

*1.* Wash TEM grid 3x by touching the grid to a drop of 100-200 ul protein dilution buffer, on the 3rd drop release the TEM grid so it floats on its surface, then pick up with Vitrobot forceps as shown on the image below. Do not submerge grids.

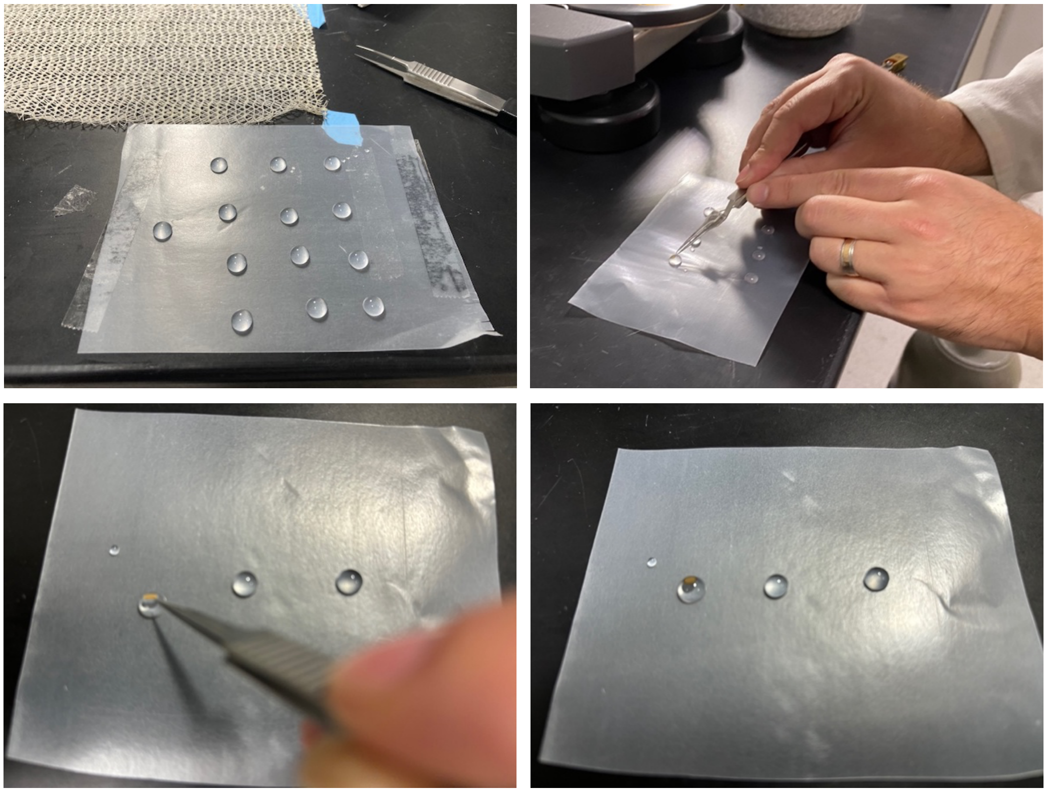

*Figure 13 - sequential steps of grid washing. Prepare a set of 100-200 uL drops, then wash the grid in three sequential drops. Leave the grid floating, but not submerged, on the last drop.*

### Ni-NTA Affinity Grid Protocol

*We recommend using a chemwipe to soak up the drops after one use to avoid cross contamination.*

2. Transfer the Vitrobot forceps onto the Vitrobot rod, in this image the lipid film (carbon side) is facing the right sample port of the Vitrobot.

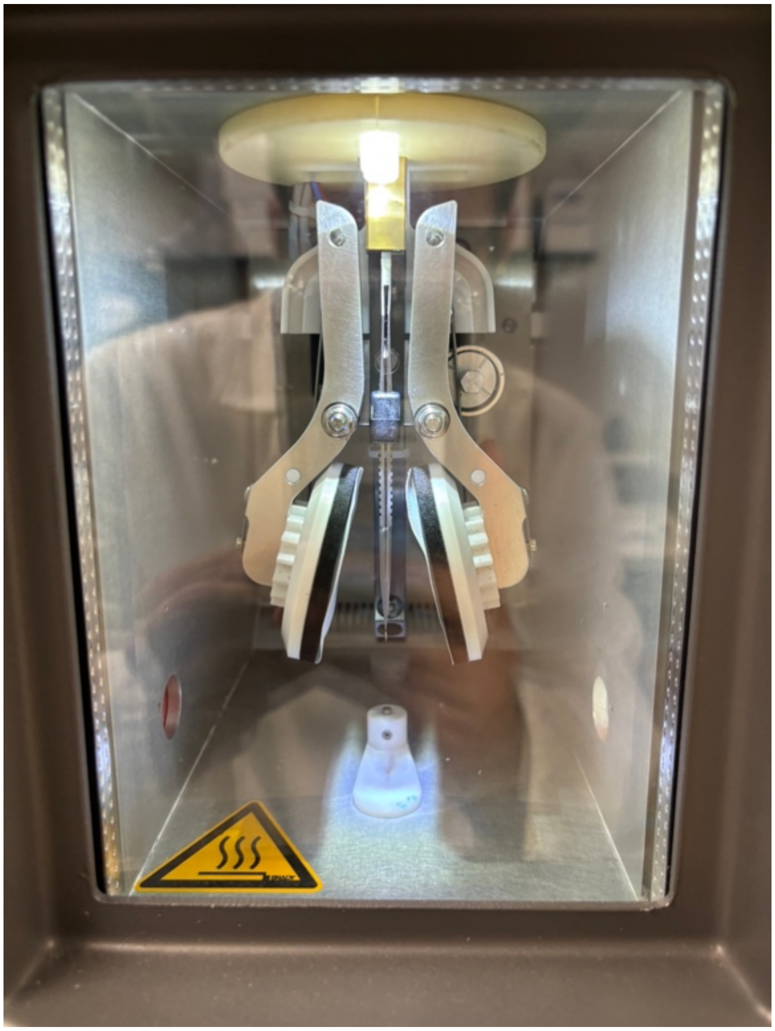

*Figure 14 - Vitrobot forceps with affinity grid inside Vitrobot humidity chamber*

3. Blot and plunge freeze into liquid ethane:propane mixture with the Vitrobot. Repeat steps to make multiple cryo-grids. We typically prepare up to 8 in one experiment. This normally takes us 4-6 hours from initial lipid coating through vitrification. Store grids under liquid nitrogen or image with the cryoTEM.
